# scAAVengr: Single-cell transcriptome-based quantification of engineered AAVs in non-human primate retina

**DOI:** 10.1101/2020.10.01.323196

**Authors:** Bilge E. Öztürk, Molly E. Johnson, Michael Kleyman, Serhan Turunç, Jing He, Sara Jabalameli, Zhouhuan Xi, Meike Visel, Valérie L. Dufour, Simone Iwabe, Felipe Pompeo Marinho, Gustavo D. Aguirre, José-Alain Sahel, David V. Schaffer, Andreas R. Pfenning, John G. Flannery, William A. Beltran, William R. Stauffer, Leah C. Byrne

## Abstract

Adeno-associated virus (AAV)-mediated gene therapies are rapidly advancing to the clinic, and AAV engineering has resulted in vectors with increased ability to deliver therapeutic genes. Although the choice of vector is critical, quantitative comparison of AAVs, especially in large animals, remains challenging. Here, we developed an efficient single-cell AAV engineering pipeline (scAAVengr) to quantify efficiency of AAV-mediated gene expression across all cell types. scAAVengr allows for definitive, head-to-head comparison of vectors in the same animal. To demonstrate proof-of-concept for the scAAVengr workflow, we quantified – with cell-type resolution – the abilities of naturally occurring and newly engineered AAVs to mediate gene expression in primate retina following intravitreal injection. A top performing variant, K912, was used to deliver SaCas9 and edit the rhodopsin gene in macaque retina, resulting in editing efficiency similar to infection rates detected by the scAAVengr workflow. These results validate scAAVengr as a powerful method for development of AAV vectors.

## Introduction

Gene therapy is a rapidly developing approach for the treatment of inherited disease, and AAV is a leading viral vector candidate for safe and efficient delivery. A growing number of clinical trials are using AAV to treat diseases such as retinal degeneration, neurological disorders, and hemophilia, through gene replacement, genome editing, and optogenetics (*1, 2*). However, significant hurdles prevent the successful, widespread implementation of AAV-mediated gene therapies, including efficient gene delivery and immune response to viral vectors and gene products. Recent efforts to reengineer viral vectors have shown promise for addressing these issues, resulting in AAVs with improved abilities(*3*). However, validation and quantitative comparison of new viral vectors remains a challenging and burdensome process. The selection of an optimal vector is essential to the success of the therapy. Sufficient gene expression is critical, while greater efficiency of gene delivery reduces the titer of vector required and decreases the likelihood of immune response. Quantitative comparisons of newly engineered vectors, including evaluation of transgene expression levels and cell-type tropism, have in the past required large numbers of animals, and therefore involved significant ethical and financial burden. Here, we have developed a single cell RNA-seq AAV engineering (scAAVengr) pipeline for rapid, quantitative *in vivo* comparison of transgene expression from newly engineered AAV capsid variants across all different cell types in a tissue in parallel, and in the same animals.

The scAAVengr pipeline was implemented in primates, which are the most physiologically similar animal to humans, and are thus a critical preclinical model. Successful clinical translation of gene therapies depends on highly efficient vectors for human tissue, and vector performance in small animals often does not accurately predict efficiency in primates. For retinal gene therapy in particular, primates are essential, as existing AAV vectors infect the primate retina significantly less efficiently than in rodent retina(*4*). Furthermore, primates are the only animal model that has a macula and foveal pit (the region of the retina responsible for high acuity vision in humans), making them the most relevant translational model. Notably, the pattern of AAV expression also differs in foveal and in peripheral retina(*4*), with highest expression in the foveola and in a perifoveal ring of retinal ganglion cells, and punctate expression near blood vessels in the periphery. We therefore developed this NHP single-cell RNA-Seq pipeline to quantitatively evaluate the clinical potential of multiple lead candidates across all retinal cell types, in the foveal and peripheral retina, in a large animal model with eyes similar to humans.

## Results

### Directed evolution of AAV vectors in canine retina

We first created additional AAV vectors with an enhanced capacity to target the outer retina following intravitreal injection, by implementing directed evolution (DE) of AAV in canines (Fig. S1–S3, and methods). DE, which involves applying a selective pressure to libraries of mutated AAV vectors, and conducting iterative rounds of selection, has been used in mouse to create AAV vectors with new abilities to infect Müller glia(*5*), to infect photoreceptors(*4*) and in primates to deliver genes to the outer retina(*6*). Here, we used DE to engineer new AAV vectors with the ability to bypass structural barriers and infect retinal cells following intravitreal injection in canine retinas. Canines are the main preclinical large animal model for development of retinal gene therapies, including the landmark gene therapy clinical trials for *RPE65*-LCA2, due to similar ocular structure and availability of homologous mutant retinal degeneration strains(*7, 8*). Therefore, we hypothesized that canines were a promising model in which to conduct a DE screen.

DE was implemented similarly to the screen previously reported in primate retina (Fig. S1 (*6*). AAV2-based libraries, including a ~588 peptide insertion library(*9*), an AAV2-Loopswap library(*10*) and an AAV2-ErrorProne library(*11*) were pooled and intravitreally injected into canine eyes. Promising variants were identified from the DE screen, based on the fold increase over the five rounds of selection, normalized to their frequencies in the starting plasmid library. Then, a secondary round of screening was performed to compare 20 top candidate canine DE variants. These 20 top vectors, along with an AAV2 control, were packaged individually with a ubiquitous CAG promoter driving expression of GFP fused to a unique DNA barcode. Vectors were titer matched, mixed together, and injected intravitreally into both eyes of 3 WT dogs (Fig. S2). Following DNA and mRNA extraction, AAV genome barcodes were PCR amplified from genomic DNA and from cDNA, from photoreceptors and RPE. Variants were ranked on the basis of the normalized change in frequency of their representation in the recovered genomes relative to the injected AAV library (% of total in recovered AAV library / % of total in injected library) (Figure S3). Rankings based on mRNA and DNA recovery indicated different top-performing variants.

The top ranked variant based on DNA recovery, K916, contains a 10 amino acid insertion (PAPQDTTKKA) at position ~588. The top ranked variant based on mRNA recovery, K912, contains a 10 amino acid insertion (LAPDSTTRSA) at position ~588. The convergent variant from the DE screen, that is, the variant that was most abundant at the end of the screen, K91, which was also overrepresented in the original library, contains a 10 amino acid insertion (LAHQDTTKNA) at position ~588. We have previously shown that convergent variants from DE screens are not necessarily top performers (*6*). That is, the variants with greatest fold increase during selection, rather than the most frequent final variant, is the optimal metric for identifying top-performing variants. An additional top-ranking variant based on DNA and RNA recovery, K94, ~588 LATTSQNKPA, was also chosen for further testing.

### Single-cell RNA-Seq quantification of AAV efficiency across all cell types

Once an initial set of vector candidates has been created, the relative fitness of these variants to infect different types of cells and retinal regions, and to generate abundant transgene expression must be precisely quantified and compared in order to identify optimally efficient vectors and the best candidate vectors for clinical translation. To this end, in order to evaluate the performance of AAV variants created through DE in canine retina, and to quantitatively compare their performance to previously engineered AAV variants created through DE in primate retina(*6*), as well as to tyrosine-mutated AAV vectors, we developed a scRNA-Seq based workflow (scAAVengr) (Fig. 1A). A set of 17 AAV vectors were packaged individually with GFP constructs fused to unique barcodes. Naturally occurring AAV variants included in the set were: AAV1, AAV2 (the parental serotype of the canine and primate DE variants), AAV5, AAV8, AAV9, and AAVrh10. Tyrosine and threonine-mutated versions of AAVs, which have been shown to prevent capsid degradation(*12, 13*), included in the set were AAV2-4YF, AAV2-4YFTV, AAV8-2YF, AAV9-2YF. DE variants included in the set were K91, K912, K916, K94, and primate DE variants NHP9, NHP26, and SCH/NHP26(*6*). In previous work, NHP9 has been shown to be highly fovea specific. NHP26 has been shown to bypass structural barriers in primate retina at decreased titer(*6*).

**Fig. 1.**
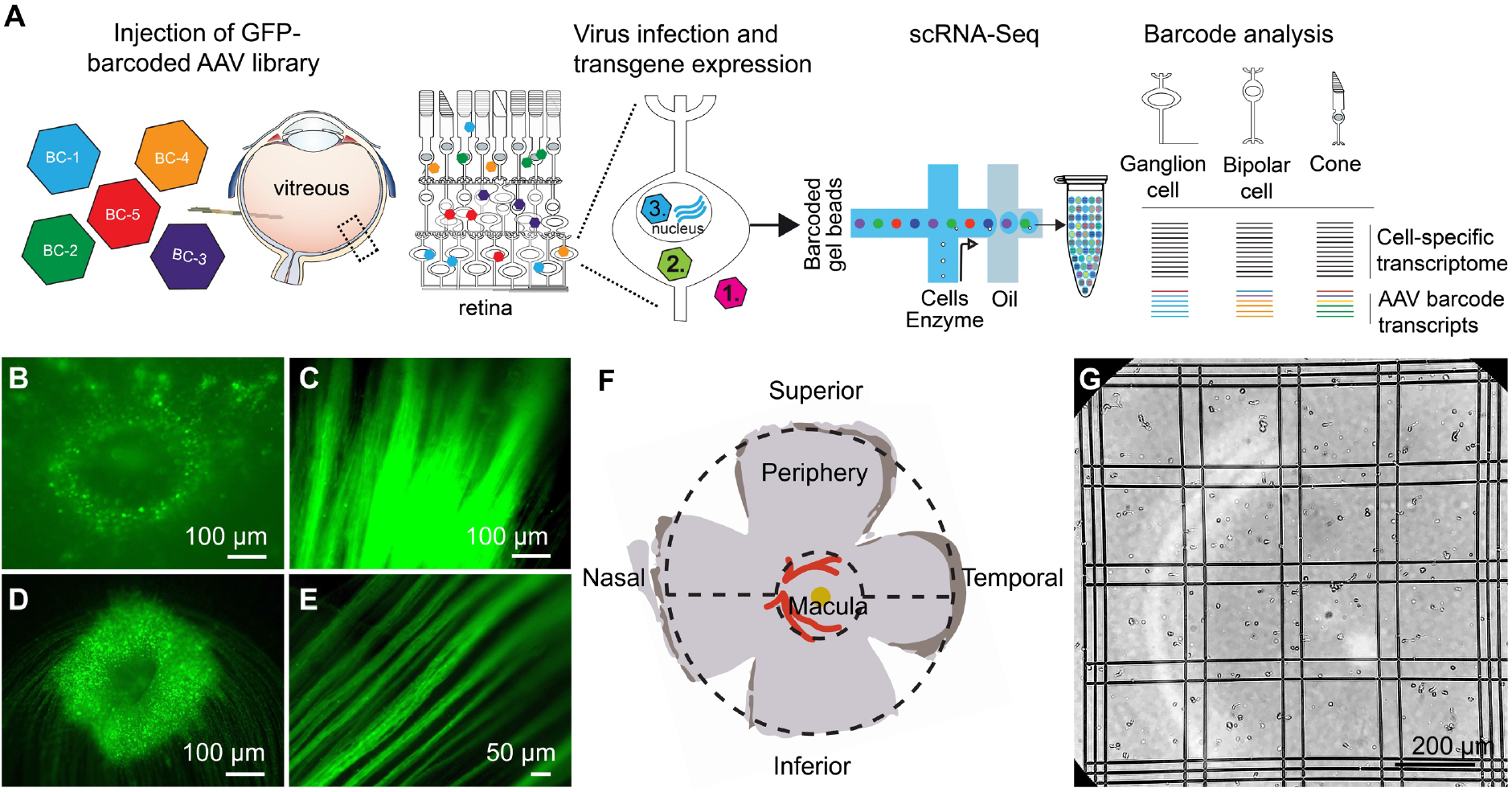
scAAVengr pipeline. **(A)** Overview of scAAVengr experimental workflow. An AAV library, consisting of variants packaged with GFP transgene fused to unique barcodes, was packaged, pooled, quantified by deep sequencing, and injected. Viruses are either noninfectious (1), bind or enter into cells but do not mediate gene expression (2), or traffic to the nucleus resulting in expression of tagged mRNA transcripts (3). Analysis took into account only viruses leading to transgene expression as in (3). Single cell suspensions of the tissue were then created, and a single cell microfluidics system was used to produce single-cell cDNA libraries. Cell types were identified by marker gene expression, and simultaneously, the ability of AAV variants to drive gene expression was evaluated based on quantification of AAV barcodes in GFP transcripts. **(B-E)** GFP-Barcoded AAV library expression in marmosets and macaques. Intravitreal injection of GFP-barcoded libraries resulted in GFP expression in the retina 8 weeks after injection. **(B)** GFP expression in the perifoveal ring in marmoset retina. **(C)** Axons from retinal ganglion cells in same injected eye as (A). **(D)** GFP expression in the perifoveal ring in macaque retina. **(E)** Axons from retinal ganglion cells in same injected eye as (c). **(F)** Diagram of primate retinal flatmount. Retinal tissue was collected from macula, and superior and inferior peripheral retina. **(G)** Retinal tissue samples were dissociated into single cell suspensions which were counted using Trypan blue. Trypan blue exclusion is also a test for cell viability. Cell suspensions were then processed through a 10X Chromium scRNA-seq controller.

Equal amounts of each GFP-barcoded virus were packaged and pooled. The representation of each variant in the packaged and pooled library was then quantified by deep sequencing. The pooled library was intravitreally injected into the eyes of 3 NHPs (2 marmosets and 1 cynomolgus macaque, Fig. 1A, and see Supplementary Table 1). Eight weeks after intravitreal injection, samples from GFP-expressing retinas were collected (Fig. 1B-E). Retinal tissue from macula and peripheral regions were dissociated into single cell suspensions (Fig 1F), a 10X microfluidics controller was used to create cDNA libraries from single cells (Fig 1G), and the cDNA libraries were then sequenced to a depth of 100,000 reads per cell. Raw sequencing reads were aligned(*14*) to the marmoset genome (Ensembl) or the cynomolgus macaque genome (UCSC) and processed with multiple QC methods: empty droplets were identified(*15*), ambient RNA was removed(*16*), doublets were removed(*17*), imputation was performed to remove the effects of sparse sampling from sequencing(*18*), single cell gene expression was normalized(*19*), and batch correction was performed(*20*). Scanpy(*21*) was used in conjunction with the Leiden clustering algorithm to assign individual cells to clusters. The hypergeometric test was then used to quantify the significance of intersection of a clusters’ differentially expressed genes with retinal cell type marker genes identified previously(*22, 23*). Each cluster was assigned a cell identity based on the most significant intersection. Clusters of all major retinal cell types were identified in marmosets and macaques, largely in agreement with previous scRNA-seq performed in primate retina(*22*). AAV barcodes were quantified using Salmon(*24*) and mapped to identified cell types using in-house scripts (Figure 2).

**Fig. 2.**
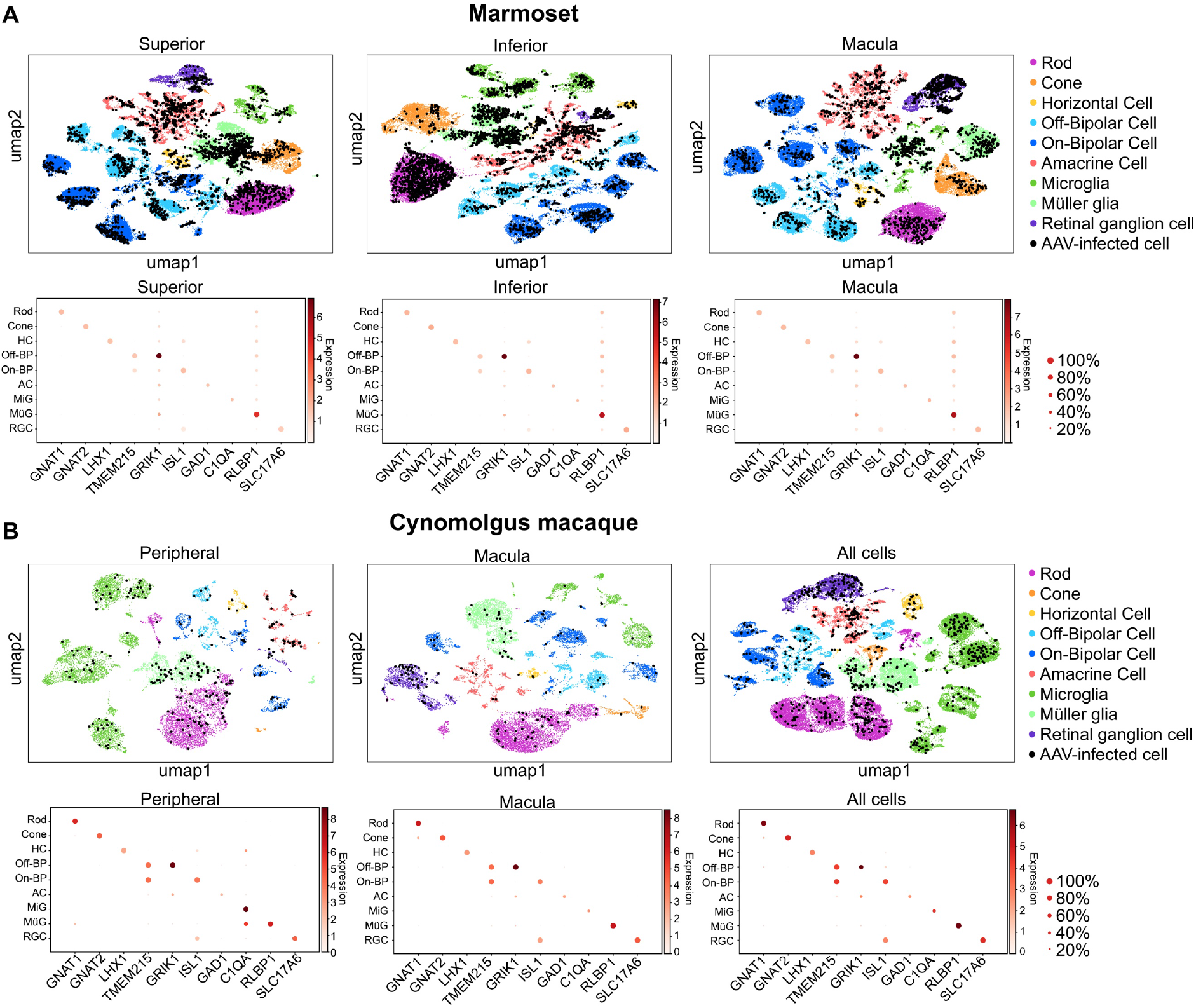
Clustering and quantification of AAV-infected retinal cells. **(A)** AAV-infected marmoset retinal cells. Maps of clustered cells from superior, inferior or macular retina show AAV infection. The cell type of each cluster is indicated by color. AAV-infected cells are shown in black. Below each cluster plot, heat maps show the marker genes used to identify cell clusters. The size of the dot indicates the percent of cells in the cluster expressing the marker gene, and the color indicates the level of marker gene expression. Data is pooled from n=2 marmosets. **(B)** AAV-infected cynomolgus macaque retinal cells. Data is from n=1 cynomolgus macaque retina, collected from peripheral or macular retina, or from the total pool of retinal cells including GFP+ FACS-sorted cells.

Three metrics were used to compare vector performance across cell types: First, the absolute number of cells infected by each serotype was quantified (Fig. S4). Second, the percent of total cells infected by each serotype was quantified for each major cell type (Fig. 3A). Third, within infected cells, the level of transgene expression was evaluated, relative to total transcripts recovered from each cell (Fig. 3B). Each of these metrics was corrected by the dilution factors for variants in the injected library, previously determined by deep sequencing. Heat maps of these metrics revealed that variants engineered through DE using canine retinas and primate retinas markedly outperformed AAV2 and AAV2 tyrosine mutants across cell types and in peripheral and macular retina.

**Fig. 3.**
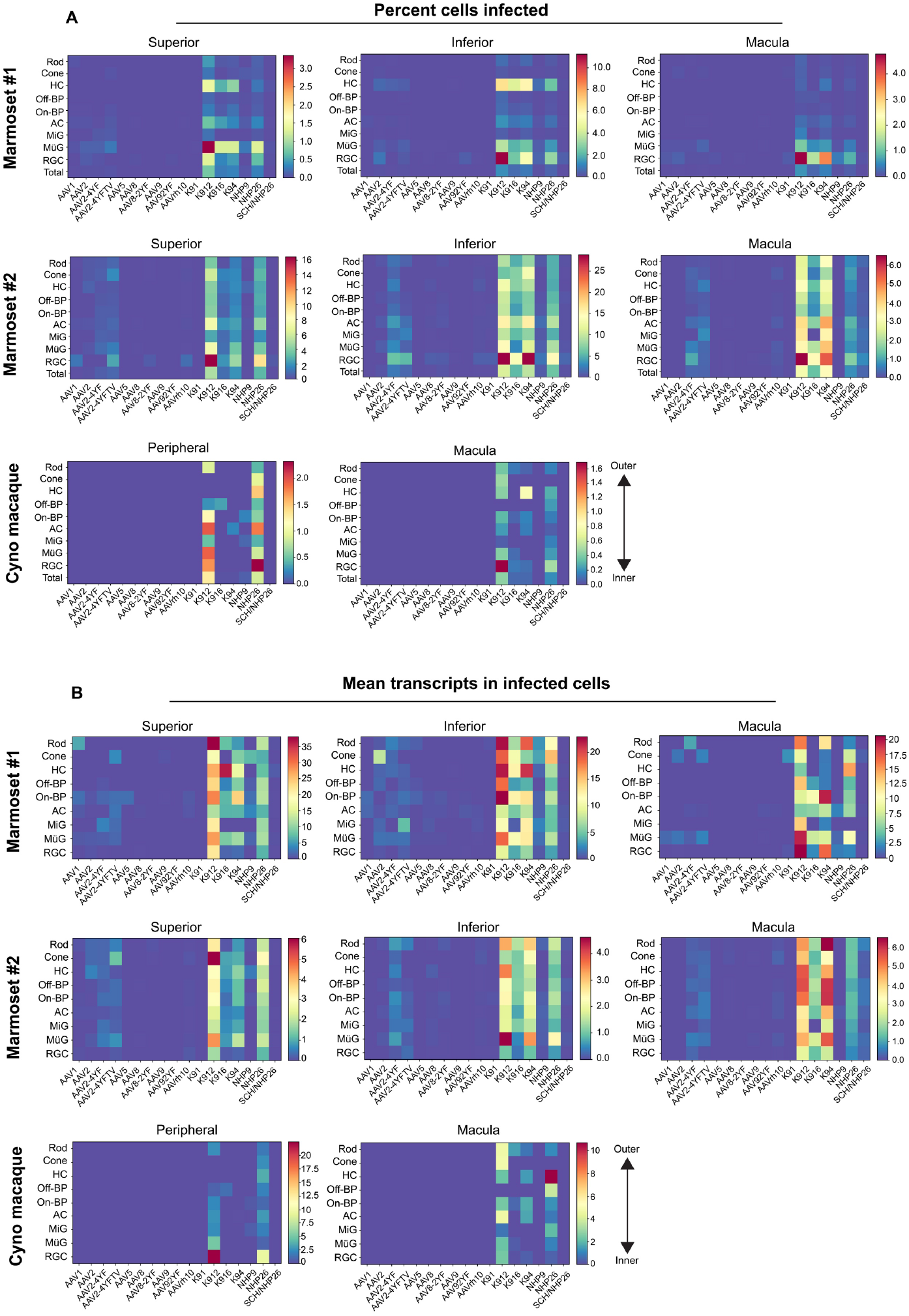
Quantitative comparison of variant infection across retinal cell types. **(A)** Percent of cells infected by AAV serotypes in marmoset and cynomolgus macaque retina. Heat maps show the percent of identified cells infected by each serotype in the screen, corrected by the AAV dilution factor, for each retinal cell type. Total = percent of all cells identified. Data is shown for each primate analyzed, across superior, inferior and macular retina. **(B)** Level of expression in infected cells. The mean level of GFP-barcode transcript expression in cells infected with AAV is shown in heatmaps, for all retinal cell types. Data is averaged across all infected cells and corrected by the AAV dilution factor. Data is shown as mean transcripts per cell/100,000 transcripts. HC = Horizontal Cell; Off-BP = Off-Bipolar Cell; On-BP = On-Bipolar cell; AC = Amacrine Cell; MiG = Microglia; MG = Müller Glia; RGC = Retinal Ganglion Cell.

Statistical analysis revealed a significant difference in the percent of total cells infected (p < 0.001, Friedman’s test, and see Supplementary Tables 2,3). Of the canine variants, K912 outperformed other engineered serotypes, in agreement with the results observed in bulk anaysis performed in dog retina (Fig. S3). The convergent variant (K91) did not outperform parental serotypes, underscoring the need for deep sequencing to determine top performing AAV variants from the DE screen. Of the primate variants, NHP26 outperformed other variants, infecting major retinal cell types in inner and outer retina in marmoset and cynomolgus macaque retina.

Evaluation of AAV infectivity at cell-type resolution revealed that newly engineered K9 variant AAVs and NHP26 infected inner and outer retinal cells in primate retina (Fig. 3). Infectivity, in terms of percent cells infected, was most efficient in RGCs and Müller glia, particularly in the macula where the inner limiting membrane is less of an anatomical barrier. In the outer retina, rods and cones were also infected. Higher rates of infection and expression levels were seen in marmosets compared to the cynomolgus macaque.

In order to rank best performing pan-retinal variants and the best performing variants by cell type, variants were plotted by the mean transcripts per cell in infected cells vs. the percent cells infected for each AAV serotype (Fig. 4). Plots were created with data from all cell types on the same plot (Fig. 4A) or in individual plots per cell type, for each region tested in each primate (Fig. 4B.) These plots revealed that K912 was the overall best performing canine variant across retinal cell types, and NHP26 was the top performing primate-derived variant across cell types. Of all the variants tested, K912 was the top performer across cell types. In order to determine the number of AAV variants infecting a single cell, upset plots were created to show the number and serotypes of AAV particles infecting individual cells. Upset plots show the number of cells infected by a particular combination of AAVs (the intersection size) as well as the number of cells infected by a particular serotype (the set size). The majority of infected cells were infected by a single variant (K912), although many cells were infected by multiple serotypes (Fig. 5). As many as 8 serotypes infected a single cell in marmoset retina, while up to 3 serotypes infected a single cell in macaque retina.

**Fig. 4.**
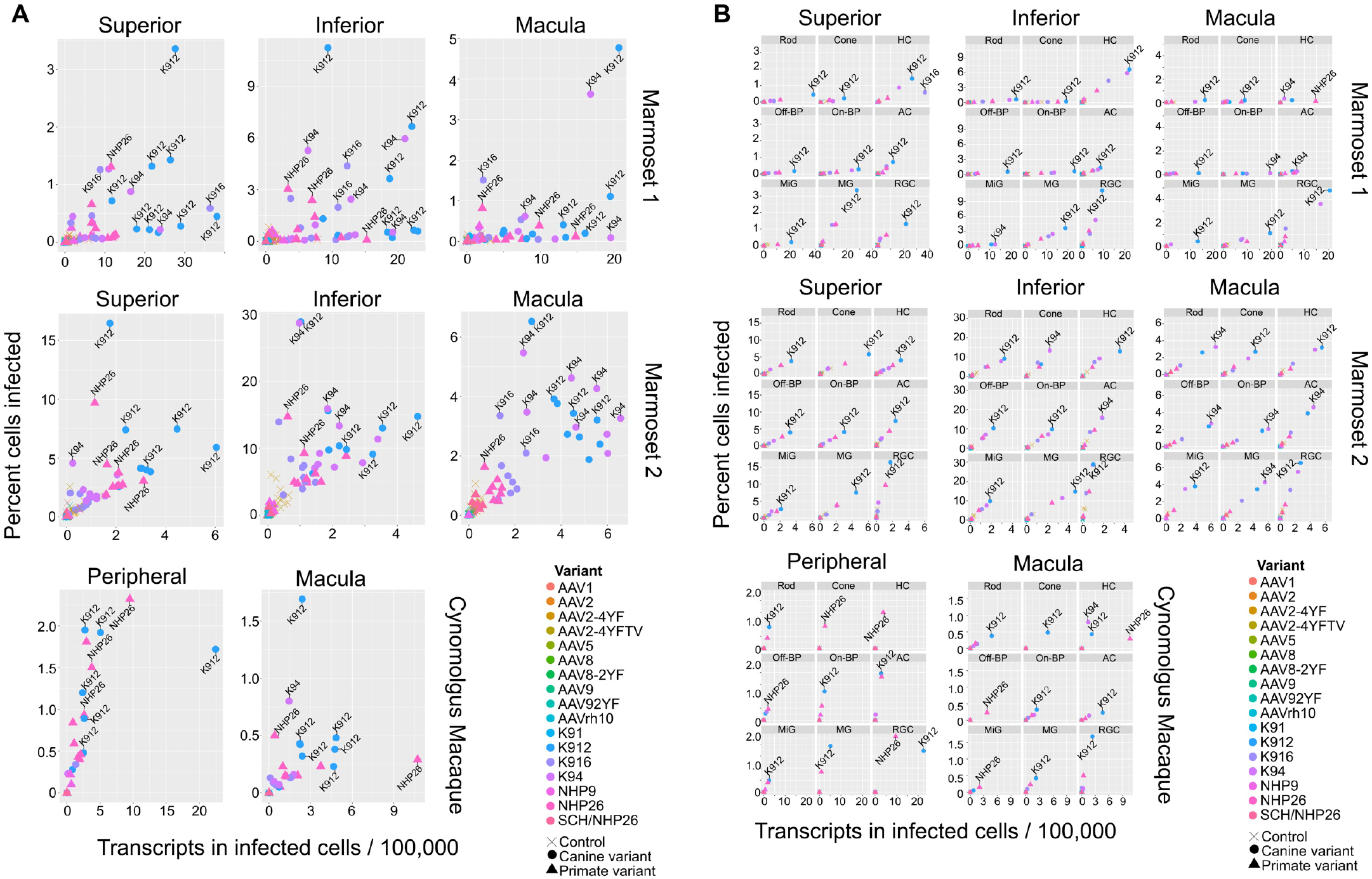
Serotype performance across retinal regions. **(A)** Scatter plots, which plot the number of transcripts in infected cells per 100,00 transcripts vs the percent of cells infected for each serotype reveal that K912 is the overall best performing canine variant across retinal regions, while NHP26 is the best performing primate DE variant. Each variant has nine data points, one for each cell type. Serotypes are indicated by color. Data is from n=2 marmosets and n=1 cynomolgus macaque. A subset of the top performing variants, according to each variable, are labeled. **(B)** AAV variant performance in each cell type. Scatter plots, which plot the number of transcripts in infected cells per 100,00 transcripts vs the percent of cells infected reveal that K912 is the overall best performing variant across most retinal cell types and across retinal regions, while NHP26 is the best performing primate-derived variant. Individual plots show the performance for AAV serotypes (in different colors) in individual cell types, across retinal regions. A subset of the top performing variants, according to each variable, are labeled. HC = Horizontal Cell; Off-BP = Off-Bipolar Cell; On-BP = On-Bipolar cell; AC = Amacrine Cell; MiG = Microglia; MG = Müller Glia; RGC = Retinal Ganglion Cell.

**Fig. 5.**
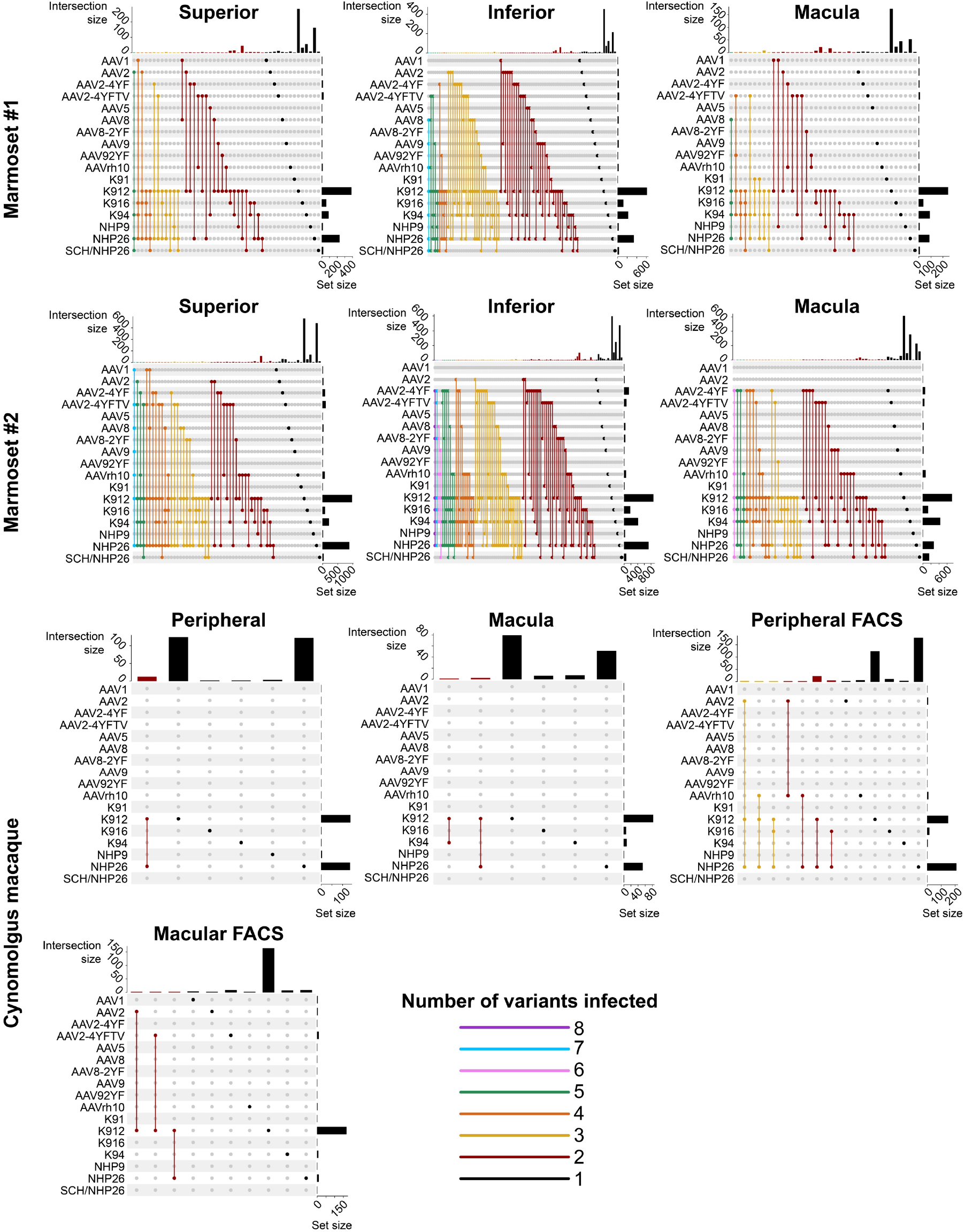
Cells infected by multiple AAV serotypes. Upset plots show that multiple AAV serotypes can infect the same retinal cell, although the majority of retinal cells are infected by the top performing variants. Plots are shown across multiple regions in marmoset and cynomolgus macaque retina. Dots and connecting vertical lines indicate the serotype and number of variants infecting single cells. The number of cells infected by a particular combination of AAVs (the intersection size) is illustrated in the bar graph across the top of the plot. The number of cells infected by a particular serotype (the set size) is shown across the right-hand Y-axis. Lines are colored by the number of AAV variants in the subset.

### Validation of K912 in primate retina

Retinal cell expression with K912, the overall top performer, was then individually validated by packaging and intravitreally injecting a self-complementary CAG-GFP construct in 2 primates (Fig. 6A-G, Fig. S5). Ten weeks after injection, GFP expression was evident in retinal flatmounts and cross sections. Confocal microscopy imaging of PNA (which labels cone inner segments)-labeled peripheral retina, imaged at the level of the photoreceptor layer, revealed GFP expression in rods and cones, which was higher in rods than in cones, in agreement with scAAVengr heat maps. Cross sections showed strong expression in RGCs and Müller glia, which was more efficient than in outer retina, particularly in the macula, also in agreement with scAAVengr heat maps.

**Fig. 6.**
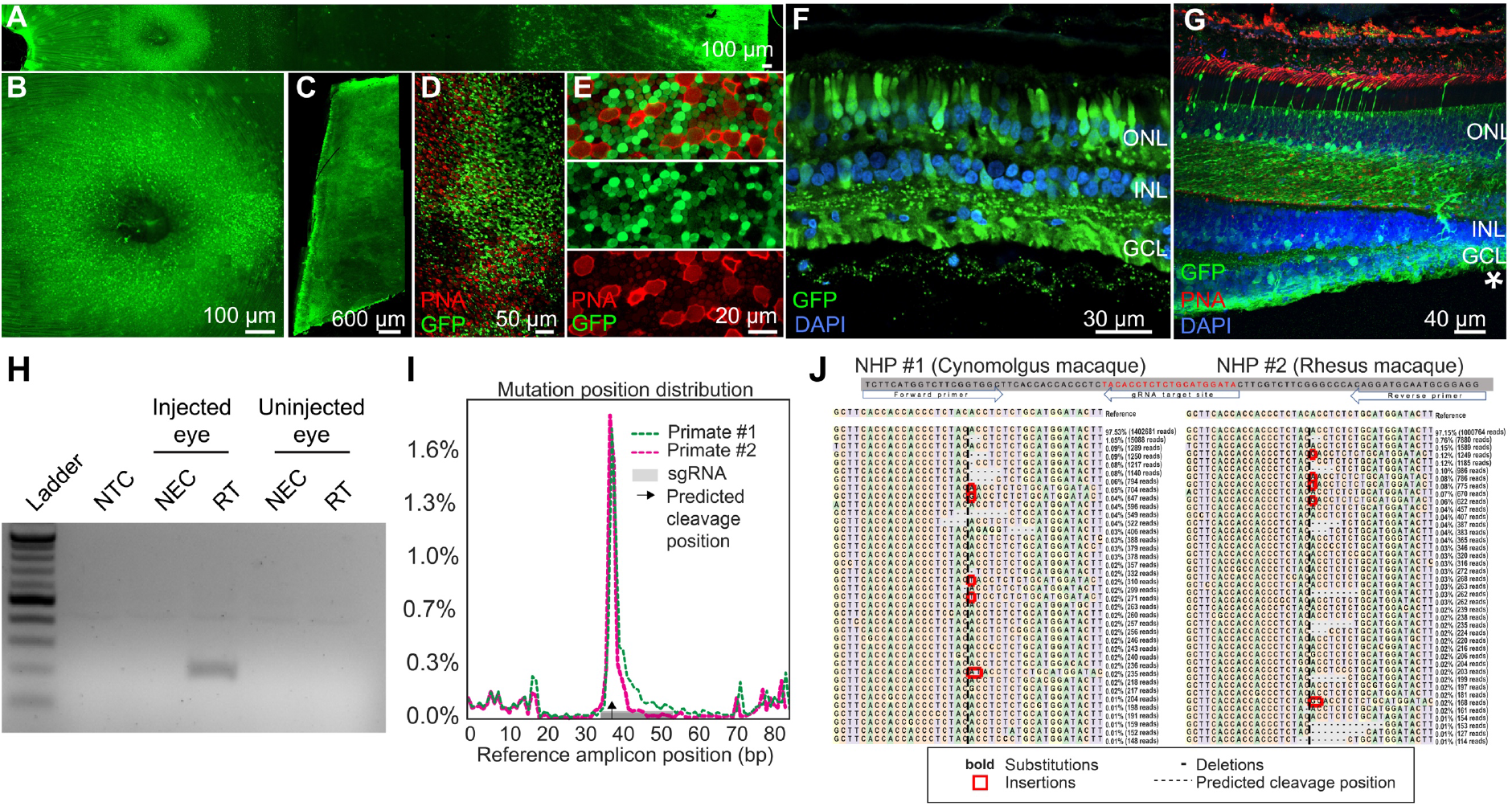
K912 expression in primate retina. **(A-G)** GFP expression in a cynomolgus macaque injected with ~2.6E+12 vg of K912-scCAG-GFP. **(A)** GFP expression in a flatmounted cynomolgus macaque retina 2.5 months after injection. **(B)** GFP expression in the perifoveal ring. **(C)** GFP expression in peripheral retina. **(D)** Flatmount imaged through the photoreceptor layer in peripheral retina showing GFP expression and PNA labeling of cones. **(E)** Higher resolution image of peripheral photoreceptors, labeled with PNA. **(F)** Cross section of peripheral retina showing GFP expression and DAPI labeling of nuclei. **(G)** Cross section through the foveal edge showing GFP expression. Cone outer segments are labeled with PNA. Nuclei are labeled with DAPI. **(H)** RT-PCR of cDNA from injected and uninjected eyes. RT-PCR shows Cas9 expression in macula of the cynomolgus macaque retina. **(I)** Percent of genome editing and location of editing relative to guide RNA sequence in 2 macaques injected with K912-scCAG-saCA9-gRNA-*RHO*. **(J)** Deep sequencing reads showing deletions, insertions and base substitutions in the cynomolgus macaque and rhesus macaque followed by injection with K912-saCas9-gRNA-*RHO*.

Then, as a functional test performed in a therapeutic context, K912 was also packaged with SaCas9 (Addgene #61591) driven by a ubiquitous CMV promoter and a guide RNA targeting rhodopsin, packaged in a single vector (Fig 6H-J). This vector was injected intravitreally in a cynomolgus macaque and a rhesus macaque. RT-PCR amplifying SaCas9 cDNA showed expression of Cas9 in injected, but not in uninjected, primate retinas (Fig. 6H). We then used deep sequencing to quantify editing from retinal punches containing all cell types, which revealed that 1.8% of reads mapping to the targeted site showed editing events in the cynomolgus macaque and 1.7% editing in the rhesus macaque (Fig. 6I,J), similar to the percent total of K912-GFP infected cells. Individual reads revealed deletions, insertions and base substitutions in the cynomolgus macaque and rhesus macaque followed by injection with K912-saCas9-gRNA-RHO.

## Discussion

Together these results validate the scAAVengr pipeline as a platform for rapid evaluation of new AAV serotypes at cell-type resolution. By simultaneously quantifying cellular and viral RNA at the single cell level, this method allows for efficient, direct, and head-to-head comparison of multiple vectors across all cell types in the same animals. This workflow enables the quantitative, head-to-head comparison of AAV vectors, directly in primates, in all cell types in parallel. Panretinal efficiency of transgene expression can be determined, as well as specificity for any cell-type of interest that can be identified by its transcriptome profile. The number and identity of unique AAV serotypes infecting a single cell can be observed, and the efficiency of a potential gene therapy can be accurately estimated.

Here we have evaluated the AAV tropism of newly engineered AAV capsid variants, created in the context of dog retina, with increased ability to infect all major retinal cell types. These variants were directly compared to variants created through DE in primate eyes, as well as naturally occurring variants and tyrosine modified versions. Further analysis could be performed, using the same dataset, to quantify infectivity in specific subtypes of retinal cell types, as subtype marker genes are validated. Additional work is required to determine the maximum number of pooled AAV variants that can be screened simultaneously though our results indicate the libraries of at least 17 variants can be evaluated. Quantification revealed that multiple AAV serotypes do infect individual cells, suggesting that competition between variants does not inhibit infection. Marmoset eyes, which are significantly smaller than human and macaque eyes, were more easily infected than cynomolgus macaques. This suggests that large primates may be of more use for accurate preclinical testing of gene delivery efficiencies. The rate of Cas9 editing in NHPs was similar to the total percent of cells infected by K912, indicating that editing was efficient in infected cells. Importantly, similar rates of infection indicate that scRNA-seq can accurately estimate the efficiency of viral gene delivery. The scAAVengr workflow is applicable to any tissue for which cell type marker genes are available, and provides a rapid, quantitative method by which AAV vectors can be rapidly evaluated for their clinical potential. This method enables the definitive ranking of AAV’s, in terms of transgene expression efficiency as well as cell-type specificity, and it will allow for optimal AAV vectors to be selected for clinical translation, improving patient outcomes.

## Materials and Methods

### Study Approval

All procedures were performed in compliance with the ARVO statement for the Use of Animals in Ophthalmic and Vision Research, and with approval for the canine studies by the University of Pennsylvania Institutional Animal Care and Use Committee (IACUC # 803813), and for the NHP studies with approval from of the Division of Laboratory Animal Resources at the University of Pittsburgh (IACUC #18042326).

#### Animals

##### Dogs

Dogs, between the age of 7-17 months, were screened for neutralizing antibodies to AAV2 as previously described(*25*). All dogs had titers <1:25. A subconjunctival injection of 4 mg of Triamcinolone Acetonide (Kenalog 40) was performed immediately after intravitreal AAV delivery. Animals were treated post-injection with daily topical application of prednisolone acetate and oral antibiotics and a tapering dose of corticosteroids. Non-invasive retinal imaging by confocal scanning laser ophthalmoscopy (Spectralis HRA+OCT, Heidelberg Engineering, Germany) was performed under general anesthesia. Overlapping *en face* images were captured using the short-wavelength (480 nm) autofluorescence imaging mode to detect GFP expression. Following euthanasia, retinal samples were collected either for DNA/RNA analysis (see above) or for immunohistochemistry after paraformaldehyde fixation and embedding. The dogs were maintained at the Retinal Disease Studies Facility, Kennett Square, Pennsylvania. Other than uveitis in 2 eyes, all other injected eyes showed no adverse events (Supplementary Table 1).

##### Primates

Marmosets, cynomolgus macaques and rhesus macaques between 3-10 years of age were used for all studies, and intravitreal injections were made with methods described previously(*4*). All NHPs used in these studies were previously screened to have anti-AAV2 neutralizing antibody titers of <1:5. Monkeys used for K912-scCAG-GFP fluorophore expression received daily oral doses of cyclosporine at a dose of 6 mg/kg for the duration of the study. Marmosets received oral daily doses of meloxicam (0.2 mg/kg) for one week after injection. At the conclusion of the experiment, euthanasia was done with an IV overdose of sodium pentobarbital (75 mg kg−1), as recommended by the Panel on Euthanasia of the American Veterinary Medical Association. A summary of minor adverse events related to the procedures is summarized in Supplementary Table 2. Other than uveitis in 2 eyes, all other injected eyes showed no adverse events.

#### AAV packaging

AAV vectors were produced in HEK293T cells (ATCC), 293AAV cells (Cell Biolabs) or AAV-293 cells (Agilent) using a double (for AAV2-7mer, LoopSwap, AAV2-ErrorProne and SCHEMA libraries) or triple transfection method(*26*). Libraries were packaged using an empirically determined molar ratio of plasmids in the packaging cell line, such that each AAV particle contained the genome encoding its own capsid(*11*). Recombinant AAVs were purified by iodixanol gradient ultracentrifugation, buffer exchanged and concentrated with Amicon Ultra-15 Centrifugal Filter Units (#UFC8100) in DPBS and titered by quantitative PCR relative to a standard curve using ITR-binding primers. The relative titer of each variant was confirmed by Illumina MiSeq sequencing.

#### Directed evolution performed in canine retina

We packaged AAV2 error prone(*11*), AAV2-7mer(*9*), loop swap(*10*) and SCHEMA libraries(*27*), which were pooled and injected intravitreally into both eyes of wild type dogs (Supplementary Fig. 1). After 6 weeks, retinas were collected, and AAV *cap* genes were recovered via PCR from retinal pigment epithelium (RPE) punches. AAV genomes were then repackaged and reinjected. Five rounds of selection were performed (Supplementary Table 1), with error prone PCR done following the third round of selection to introduce additional diversity into the library. Promising variants were then identified from the screen, based on the fold increase over the five rounds of selection, normalized to their frequencies in the starting plasmid library. Twenty top variants with the largest fold increases during the overall selection were then chosen for a head-to-head analysis in canine retina. To compare the selected variants head-to-head, these 20 vectors, along with an AAV2 control, were packaged individually with a ubiquitous CAG promoter driving expression of GFP fused to a unique DNA barcode. Vectors were titer matched, mixed together, and injected intravitreally into both eyes of 3 WT dogs. Six weeks after injection, potent GFP fluorescence was detected by fundus imaging of the canine eyes (Fig. 1d, and Supplementary Fig. 2). Eyes were harvested, tissue samples were collected from across the retina (Fig. 1e), the RPE was separated from the neuroretina, and photoreceptors were collected using transverse sectioning on a cryostat(*6*). GFP expression was present in every layer of the dog retina (Fig. 1f). Following DNA and mRNA extraction, barcodes were PCR amplified from genomic DNA and from cDNA, from photoreceptors and RPE.

GFP barcodes amplified from the outer nuclear layer (ONL), RPE, and the injected libraries were then subjected to Illumina sequencing to quantify the representation of each of the variants. Variants were ranked on the basis of the normalized change in frequency of their representation in the recovered genomes relative to the injected AAV library (% of total in recovered AAV library / % of total in injected library) (Table 1). Selected variants largely outperformed AAV2 (magenta) in 3 dogs and across peripheral, mid-peripheral and central retina (Supplementary Fig. 2). In addition, the most abundant variant (K91, LAHQDTTKNA, in green), which was overly represented in the original library, did not outperform other variants in canine retina, indicating that the metric of quantity of representation in the final round of selection is not the best indicator of fitness for transgene expression. Rankings based on mRNA and DNA recovery indicated different top-performing variants. Evaluation on the basis of mRNA is a more relevant readout of AAV performance, as it is indicative of transgene expression, rather than persistence in extracellular spaces of the tissue or viral endocytosis without useful mRNA expression.

#### AAV capsid variant selection in dog neuroretina

Intravitreal injections (150 – 250 μL) were performed with a 30-gauge insulin syringe under general anesthesia delivering the viral solution in the mid-vitreous. Three weeks later, dogs were euthanized by intravenous injection of sodium pentobarbital, and both eyes were flattened by making relief cuts in the globe. 2 mm punches of neuroretina and RPE were immediately collected from superior, inferior, temporal, and nasal regions of the retina, as well as from the area centralis, and flash frozen. DNA was extracted from samples using a Qiagen DNeasy blood and tissue kit, according to the manufacturer’s instructions.

#### Deep sequencing of AAV libraries from rounds of selection conducted in dogs

A ~75-85 base pair region containing the 7mer insertion was PCR amplified from harvested DNA. Primers included Illumina adapter sequences containing unique barcodes to allow for multiplexing of amplicons from multiple rounds of selection (Supplementary Table 7). PCR amplicons were purified and sequenced with a 100-cycle single-read run on an Illumina HiSeq 2500. DNA sequences were translated into amino acid sequences, and the number of reads containing unique 7mer insert sequences were counted. Read counts were normalized by the total number of reads in the run. Pandas was used to create plots.

#### Deep sequencing analysis of rounds of selection in canine

Best performing variants were chosen as variants with the greatest fold increase in the final round of selection relative to the initial plasmid library (# reads in final round, normalized to total number of reads in the round / # of reads in plasmid library, normalized to total number of reads in the round). A pseudo-count of 1 was added to each variant in every round, in order to mitigate effects of small number increases and allow analysis of variants with a zero count in sequencing of the original library(*28*).

#### GFP barcoded AAV library construction

Unique 25 bp DNA barcodes were cloned after the stop codon of eGFP, in an AAV ITR-containing plasmid construct containing a self-complementary CAG promoter driving eGFP expression (scCAG-eGFP-Barcode-bghPolyA). Individual AAV variants were packaged separately with constructs containing different barcodes using a triple transfection method. Variants were then titer matched and mixed in equal ratios before injection into dogs, and primates.

#### Deep sequencing of GFP-barcode AAV library screened in dogs

DNA and mRNA were extracted from retinal samples using a Qiagen Allprep kit. Samples were collected from areas across the retina, and from ONL or RPE. cDNA was created from mRNA using Superscript III reverse transcriptase, according to the manufacturer’s recommendations. Barcodes were PCR amplified directly from DNA or cDNA. Primers amplified a ~50 bp region surrounding the GFP barcode and contained Illumina adapter sequences and secondary barcodes to allow for multiplexing of multiple samples (Supplementary Table 7). PCR amplicons were purified and sequenced with a 100-cycle single-read run on a MiSeq. Read counts were normalized by total number of reads in the run. Analysis of barcode abundance was performed using in-house code written in Python, followed by creation of plots in Pandas. Best performing variants were selected based on the fold increase in the percent of total library, relative to the injected library (% of total in recovered sample / % of total in injected library). Analysis was performed on n=3 dogs.

#### Primers

Primer sequences are listed in Supplementary Table 4.

#### Histology in primates

For histology, both retinas were lightly fixed in 4% paraformaldehyde, and transferred to PBS. Retinas were then embedded in 5% agarose and sectioned at 100 μm on a vibratome. Tissue was then examined by confocal microscopy. Antibodies for labeling were anti-GFP (A11122, Thermo, 1:250) and peanut agglutinin (PNA, Molecular Probes, 1:200) a lectin that specifically binds to the cone photoreceptor extracellular matrix.

#### Pooling and quantification of GFP-barcode libraries in primates

Equal quantities of AAV serotypes were packaged separately and pooled. Deep sequencing was used to quantify the relative abundance of vector in the pooled library ~2.65E+11 – 6.58E+11 vg/ml), by amplifying using primer/adapters and sequencing on a MiSeq Nanoflow cell. Dilution factors were determined to be: AAV9: 1, AAVrh10: 0.62, AAV8: 0.54, SCH-NHP9/26: 0.46, NHP26: 0.31, AAV9-2YF: 0.22, AAV8-2YF: 0.21, K912: 0.21, AAV2-4YF: 0.17, K91: 0.16, AAV2-4YFTV: 0.13, AAV5: 0.11, K94: 0.11, K916: 0.08, AAV2: 0.08, NHP9: 0.07, AAV1: 0.07.

#### Single-cell dissociation of primate retina

The NHP retinas were dissected and regions of interest were isolated (macula, superior and inferior periphery). For cynomolgus macaque, superior and inferior periphery were pooled. Retinal tissue was placed in Hibernate solution (Hibernate A -Ca Solution, BrainBits LLC), and cells were then dissociated using Macs Miltenyi Biotec Neural Tissue Dissociation Kit for postnatal neurons (130-094-802) according to manufacturer’s recommendations. Dissected retina pieces were incubated with agitation at 37 °C and further mechanically dissociated. The dissociated neural retina was filtered using a 70 μm MACS Smart Strainer (Miltenyi Biotec) to ensure single-cell suspension. Cells were resuspended in 0.1% BSA in D-PBS and processed immediately for scRNA-seq.

#### FACS

In the rhesus macaque, after dissociating cells using Macs Miltenyi Biotec Neural Tissue Dissociation Kit for postnatal neurons (130-094-802), a Miltenyi MACS Tyto sorter was used to enrich for GFP-positive cells. Cells were resuspended 0.1% BSA in D-PBS and processed immediately for scRNA-seq.

#### Single cell RNA-seq

Marmoset and cynomolgus macaque samples were prepared for single cell analysis using a 10x Chromium Single Cell 3’ v3 kit. Briefly, single cells from retina samples were captured using 10X Chromium system (10X Genomics), the cells were partitioned into Gel beads-in-emulsion (GEMS), mRNAs were reverse transcribed and cDNAs with 10X Genomics Barcodes were created with unique molecular identifiers (UMIs) for different transcripts. Purified cDNA was PCR amplified and further purified with SPRIselect reagent (Beckman Coulter, B23318). Final libraries were generated after fragmentation, end repair, A-tailing, adaptor ligation, and sample index PCR steps according to 10x Single Cell 3’ workflow. An additional targeted sequencing analysis was run on these 10x-prepped cDNA samples, using PCR amplification with Q5 High Fidelity DNA Polymerase to target the GFP sequence and its associated AAV barcode. Libraries were pooled and all samples were submitted for deep sequencing on an Illumina Novaseq S4 flowcell at the UPMC Genome Center. Sequencing depth was targeted at 100,000 reads per sample for the standard scRNA-seq analysis.

##### Single cell RNA-seq pre-processing

Sequencing data was demultiplexed into sample-level fastq files using Cell Ranger mkfastq (v3 10X Genomics). Alignment and cell demultiplexing were run using STARsolo(*14*) (v2.7) with default parameters. DropletUtils(*15*) (v1.4.3) was used after STARsolo to remove empty droplets (lower.prop=0.05). Cynomolgus macaque samples were aligned to the Macaca_fascicularis_5.0/macFas5 reference obtained from UCSC and marmoset samples were aligned to ASM275486v1 obtained from Ensembl. Gene annotation for the cynomolgus macaque was created by lifting over the pre-mRNA gene annotations from the hg38 Ensembl human genome. ASM275486v1 gene annotation files from Ensembl were used for the marmoset.

Cell-free RNA contamination in droplets was estimated using SoupX(*16*) (v0.3.1). We estimated contamination using genes selected from SoupX’s inferNonExpressedGenes method, which identifies genes with highly bimodal expression in the samples. The gene expression in cynomolgus macaque samples was adjusted according to the SoupX estimates, using the ‘adjustCounts’ method. No indication of cell-free RNA contamination was observed in marmoset samples, based off of global expression of key marker genes, and therefore gene expression was not adjusted.

Doublets (10x droplets containing two cells instead of one) were then identified using SCDS(*17*) (v1.0.0). Any droplets with a hybrid score > 1.3 were considered doublets. Size factor normalization of the single cell gene expression was achieved using Scran(*19*) (v1.12.1), and replicates as well as left/right eyes of the same region were combined for normalization. Finally, imputation strategies were used to denoise the high sparsity that is common in scRNA-sequencing (ALRA v1.0(*18*)).

##### Single cell RNA-seq cell identification

Scanpy(*21*) (v1.4.4.post1) was used for the analysis of the scRNA-seq data. First, the top 50 principal components of the gene expression matrix were computed and the Euclidean distance between cells was calculated in this low dimensional space. Then, the distances of 0.5% of the closest neighbors were kept for each cell and embedded into a neighborhood graph using the UMAP algorithm. Finally, Leiden clustering was performed on the single cell neighborhood graph. Batch correction was performed to combine samples within the same species (including samples across the two marmosets as well as FACS-sorted/non FACS-sorted cynomolgus macaque samples) using Scanorama(*20*) (v1.2) and clustering was performed on the batch-corrected values. If samples were batch corrected, normalized counts were saved as raw data and used for differential gene expression analysis.

Cell types were determined by running a differential gene expression analysis using Scanpy’s “rank_gene_groups” function. We used a hypergeometric test and calculated the significance of the intersection of marker genes from one cluster with the published marker genes of each retinal cell type. A Bonferroni p-value correction was applied to account for multiple-hypothesis test. Each cluster is assigned a cell type based on the most significant marker gene intersection p-value. For clusters where the hypergeometric test could not identify a specific cell type match, we annotated the cell type based on marker gene expression using a known cell type marker database. We used two scRNA-seq retina papers(*22, 23*) to construct our database of marker genes as well as a larger aggregated scRNA-seq marker database(*29*).

### Statistical Analysis

Statistical tests were run in R. Normality of the datasets was checked using the Shapiro-Wilk test, and it was found that the datasets were unlikely to be normally distributed (p-values < 2.2E-16 for percent cells infected, p-values < 5.86E-13 for mean transcripts in infected cells). Friedman’s test was run on 8 samples from the marmoset and cynomolgus macaque comparing the total percent of all cells infected in each sample across the AAV variants. Additional Friedman’s tests were run for each cell type, analyzing the percentage of cells infected across variants on a cell type level. One-sided Wilcoxon signed-rank tests were run on the same datasets (total cells and individual cell types), comparing K912 or NHP26 with the other AAV variants, and the Benjamini-Hochberg method was used to correct p-values. AAV variants were also compared by analyzing the average transcripts in infected cells using the same statistical procedure (Supplementary Tables 3-6).

### PCR amplification of AAV barcodes for scAAVengr analysis

The performance of AAV variants is analyzed based on quantification of AAV variant-mediated GFP-barcode mRNA expression. GFP-barcodes were analyzed from (1) the original scRNA-seq data and (2) PCR amplification of GFP from the 10x single cell prepped sample library. In both cases, the GFP-barcodes were identified using Salmon(*24*) (v0.9.1) transcript quantification. Only reads with 1 hit to a GFP barcode were kept. Using these reads, AAV variants were identified based on the GFP barcode. 10x barcodes in the reads from the PCR amplification analysis were corrected according to the 10x Cell Ranger count algorithm to mitigate any errors that may have been introduced by multiple rounds of PCR. Rarely, multiple AAV variants were found per UMI (unique molecular identifier), and in this case the AAV variant with the highest number of counts for that UMI was kept.

The PCR-amplified barcodes resulted in a higher number of AAV variants found and 10x barcodes with AAV that were identified from the original scRNA-seq data were added to this set. AAV variants from the scRNA-seq dataset were added to the set if that 10x barcode was present in the PCR-amplified analysis but that AAV variant was not previously reported.

After identifying AAV variants for each 10x barcode, the 10x barcodes were mapped to the cell types identified previously during the standard scRNA-seq analysis. Once mapped to their respective cell types, AAV counts were normalized by dividing by the total transcriptome nUMI for that 10x barcode and corrected by the dilution factor for each AAV variant. Variants were then divided by the dilution factor of the variant with the highest percentage of cells infected.

### CRISPR-Cas9 editing analysis

K912 was packaged with an SaCas9 construct (Addgene; pX601-AAV-CMV::NLS-SaCas9-NLS-3xHA-bGHpA;U6::BsaI-sgRNA (Plasmid #61591). The gRNA was designed to target 285bp downstream of the *RHO* start codon. A cynomolgus macaque and a rhesus macaque were both injected intravitreally, and 9 weeks (cyno) or 6 weeks (rhesus) later, tissue was collected for processing. Genomic DNA was extracted using a Qiagen DNeasy Kit and the target site in *RHO* gene was PCR amplified with primers attached to Illumina adapter sequences. Amplicon sequences targeting *RHO* were sequenced on the Illumina iSeq and ~1,000,000 reads were recovered for each sample. CRISPResso2(*30*) (v2.0.34) was used to quantify and visualize the edits, using the amplicon sequence and guide sequence as input. Reads were filtered using an average base quality of 30 and single base quality of 20.

## General

Deep sequencing in canines was performed at the UC Berkeley Vincent J. Coates sequencing facility and the UPMC Genome Center. Confocal imaging of dogs was performed at the Berkeley Biological Imaging Facility. At the University of California Berkeley we thank Tim Day, Yvonne Lin, Cécile Fortuny, and Jaskiran Mann for technical advice and assistance. At the University of Pennsylvania, we thank Lydia Melnyk for research coordination, and the staff at the Retinal Diseases Study facility (University of Pennsylvania) for animal care. At the University of Pittsburgh, we thank DLAR staff for animal care. We thank Peter Strick for breeding of marmosets.

## Author contributions

BEO: Conceived, planned and executed experiments. Analyzed data. Wrote the manuscript. MEJ: Conceived, planned and executed experiments, analyzed data. Wrote the manuscript. MK: Analyzed data. Wrote the manuscript. ST: Performed experiments, Wrote the manuscript. JH: Performed experiments. Wrote the manuscript. SJ: Performed experiments, Wrote the manuscript. MV: Planned experiments, analyzed data. Wrote the manuscript. ZX: Performed experiments. Wrote the manuscript. VLD: Executed *in vivo* studies in dogs and analyzed data. FPM: Executed *in vivo* studies in dogs and analyzed data. SI: Executed *in vivo* studies in dogs and analyzed data. GDA: Analyzed data from *in vivo* studies in dogs. Wrote the manuscript. Wrote the manuscript. DVS: Conceived, planned and supervised execution of directed evolution performed in dogs. Wrote the manuscript. JAS: Supervised work. Wrote the manuscript. ARP: Supervised work, analyzed data. Wrote the manuscript. JGF: Conceived, planned and supervised directed evolution of AAV in dogs. Supervised Wrote the manuscript. WAB: Conceived, planned and supervised execution of *in vivo* studies in dogs. Analyzed data and wrote the manuscript. WRS: Conceived, planned and executed experiments. Supervised work. Analyzed data. Wrote the manuscript. LCB: Conceived, planned and executed experiments. Supervised work. Analyzed data. Wrote the manuscript.

## Data availability

Data is available on Dash, the University of California data sharing service, including raw counts from deep sequencing datasets in canines (Byrne, Leah et al. (2018), Directed Evolution of AAV for Efficient Gene Delivery to Canine and Primate Retina - Raw counts of variants from deep sequencing, UC Berkeley Dash, Dataset, https://doi.org/10.6078/D1895R). Cell-by-gene matrices and fastq files from scRNA-seq analysis in primates are available upon request and will be posted on NCBI.

## Supplementary Materials

**Fig. S1.**
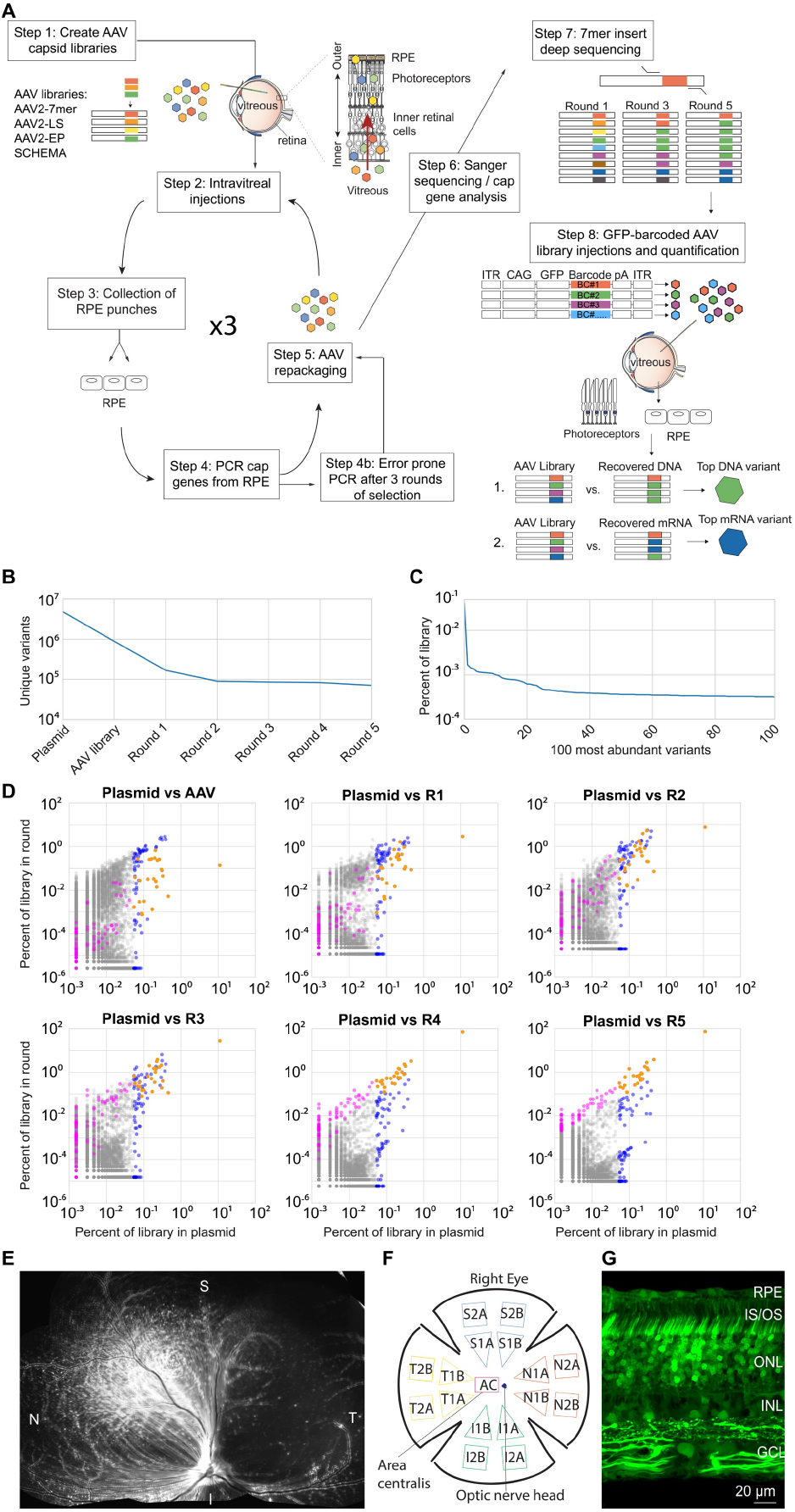
Directed evolution performed in dogs. **(A)** Highly diverse (~1E+7) libraries of AAV capsid variants were packaged such that each virus contained a genome encoding its own capsid. Libraries were pooled and injected intravitreally in canines or primates. After AAV infection had occurred, RPE punches were collected, and *cap* gene variants were PCR amplified, recloned, and repackaged for the subsequent round of injection. Five rounds of selection were performed, and error prone PCR was performed after the third round to introduce additional diversity into the library. Following the selections, each pool was subjected to deep sequencing in order to analyze the dynamics of each individual variant and overall convergence of the library. Based on their increase in representation relative to the original library, individual variant capsids were chosen and used to package a scCAG-eGFP genome also containing a unique DNA barcode sequence. These barcoded vectors were then pooled in equal amounts and injected intravitreally. Punches of neuroretina and RPE were harvested, GFP barcodes were PCR amplified from the collected tissue, and deep sequencing was used to quantify the relative abundance of barcodes. The top-performing variants were identified as those with the greatest fold increase of barcodes recovered from collected tissue relative to the injected library. **(B)** Deep sequencing revealed ~4.8E+6 variants in the AAV2-7mer library, which converged to ~7.1E+4 variants in the final round of selection. **(C)** Analysis of variant copy number in the plasmid library revealed that variants were not equally represented, with some variants having 100-fold more copies than others. **(D)** Scatterplots illustrate the behavior of individual variants through each round of selection. Each dot represents an individual variant. Variants overrepresented in the original library are colored blue. Variants with the greatest fold increase in representation in the final round of selection are colored magenta. Variants that were both overrepresented in the original library and increased significantly in representation over rounds of selection (an overlap of blue and magenta) are colored orange. **(E)** The scCAG-eGFP-barcoded AAV library composed of 20 top variants, and co-injected intravitreally into a WT dog eye, resulted in robust GFP fluorescence in the retina 3 weeks after injection. **(F)** A map illustrates the location of samples collected from the GFP-barcoded library-injected retinas. **(G)** Confocal imaging of canine retina (superior quadrant) showed GFP expression in all retinal layers after injection of GFP-barcoded AAV libraries. RPE = Retinal pigment epithelium; IS/OS = inner and outer segments; ONL = outer nuclear layer; INL = inner nuclear layer; GCL = ganglion cell layer.

**Fig. S2.**
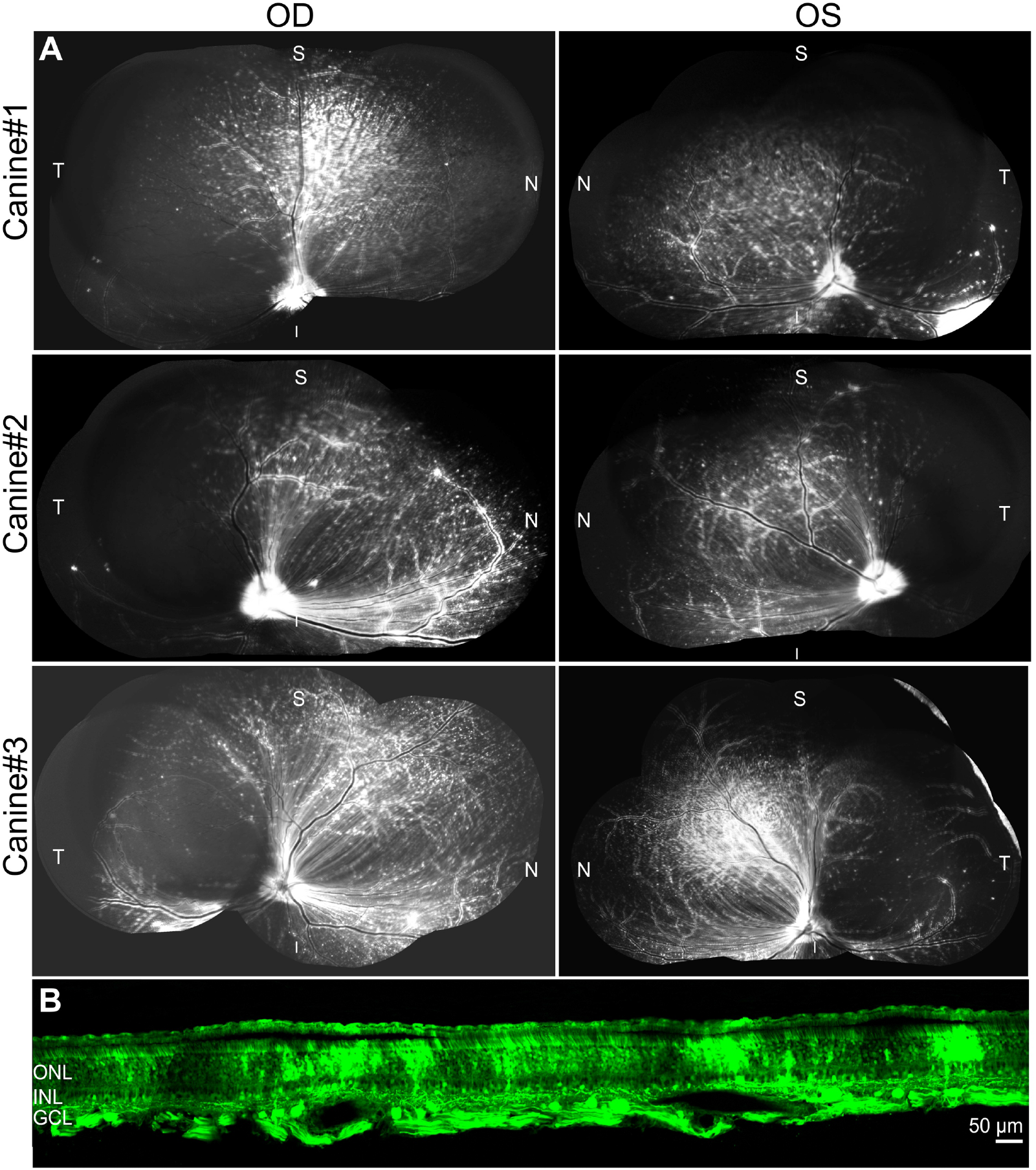
Injection of GFP-barcoded AAV library of top AAV variants in dogs. **(A)** Intravitreal injection of GFP-barcode library in 3 dogs resulted in transgene expression 3 weeks after injection. **(B)** Imaging of GFP in a cross section through the superior quadrant of the retina revealed GFP expression in all retinal layers.

**Fig. S3.**
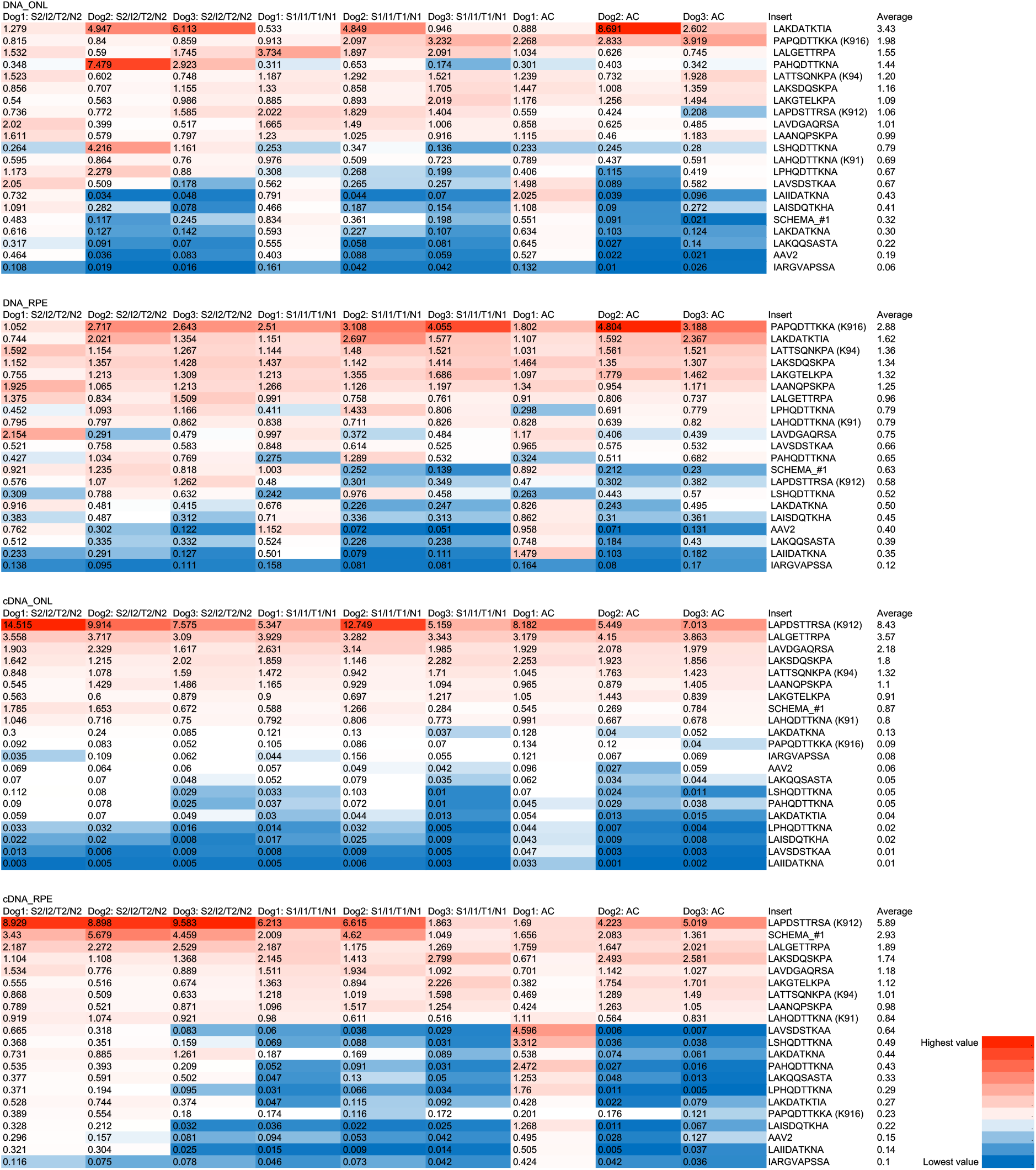
Performance of AAV variants following intravitreal injection of GFP-barcode library in dog. GFP barcodes amplified from the ONL and RPE were subjected to Illumina sequencing to quantify the representation of each of the variants. Heat maps show the performance of variants, ranked on the basis of the normalized change in frequency of their representation in the recovered genomes relative to the injected AAV library (% of total in recovered AAV library / % of total in injected library). Best performing variants recovered from DNA and mRNA are shown, across pooled samples from 3 retinal regions in 3 dogs.

**Fig. S4.**
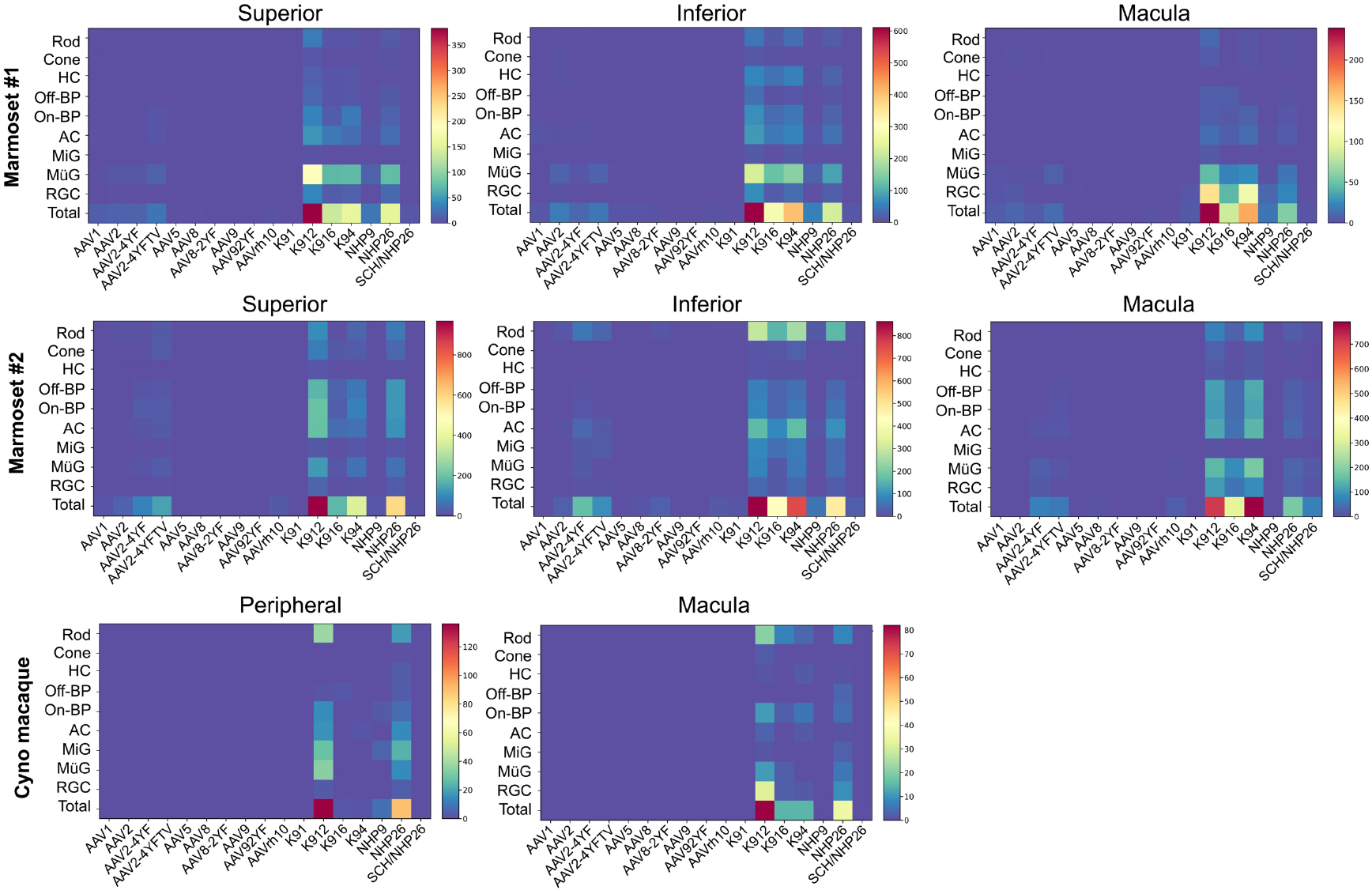
Numbers of AAV-infected cells. Heat maps show the number of cells infected by each serotype of virus, across retinal regions, in marmosets and cynomolgus macaque. Percent of cells infected is shown in Fig. 3. Numbers are corrected by dilution factor of the virus.

**Fig. S5.**
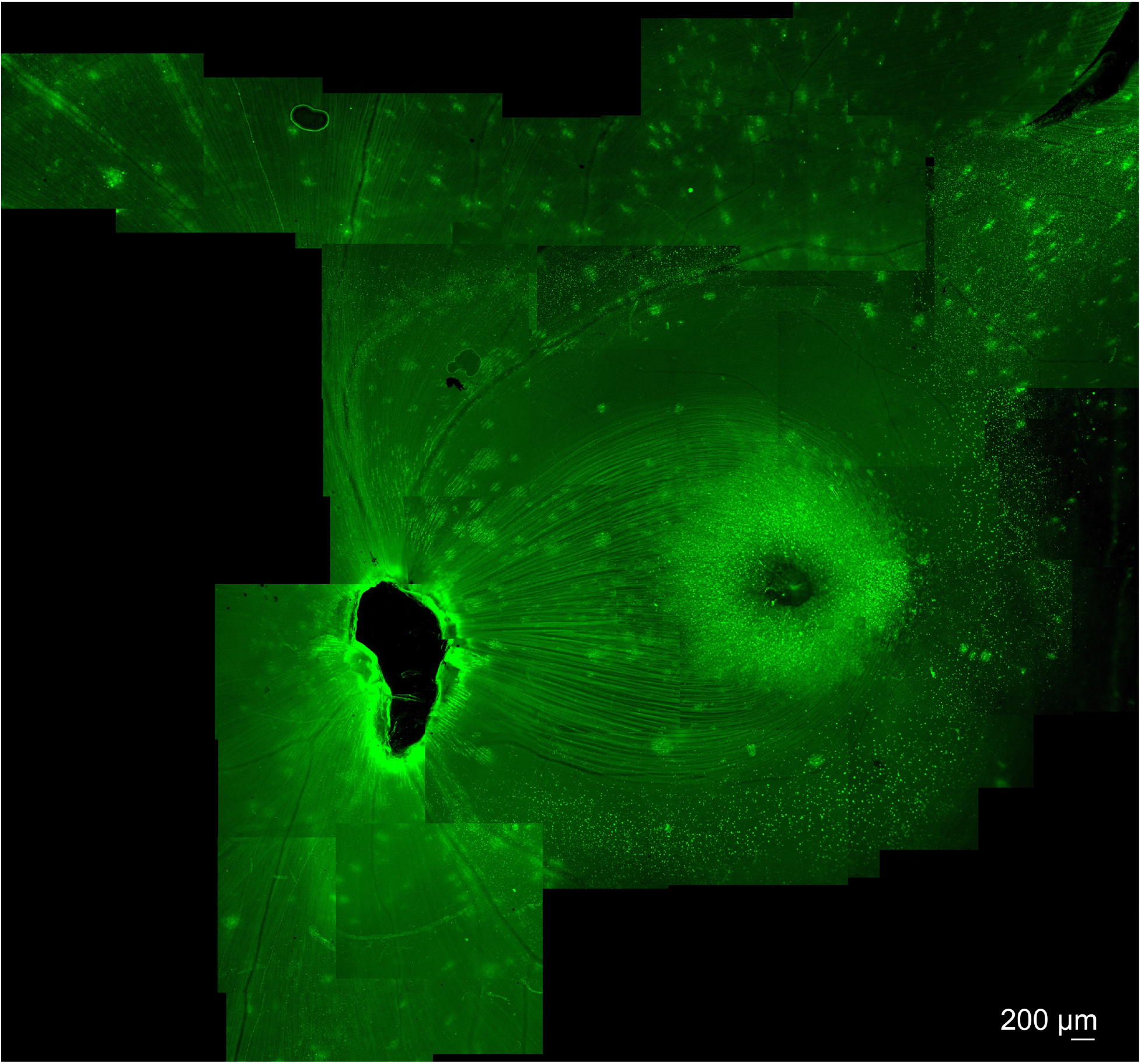
Expression of K912-CAG-GFP in cynomolgus macaque retina. Intravitreal injection of K912-scCAG-GFP results in GFP expression in the central macaque retina. Expression results in bright expression in a perifoveal ring, and in punctate regions around retinal blood vessels. Imaging was performed 10 weeks after injection.

---

**Table S1.**
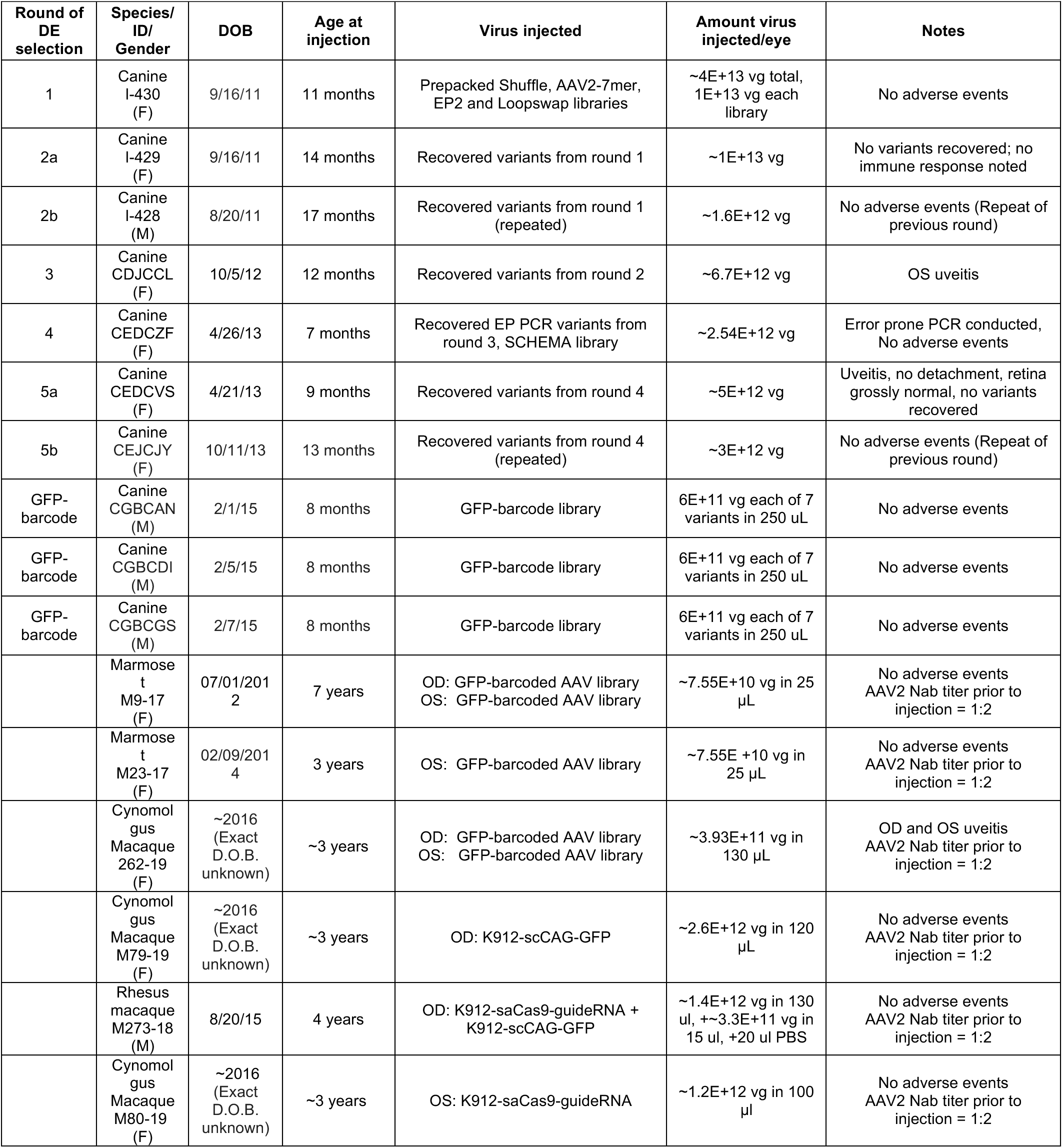
Summary of injections performed. Summary of intravitreal AAV injections performed in dogs and primates. AAV selection rounds in canines 2b and 5b were repeated selections of the previous rounds, which did not result in the amplification of AAV variants.

**Table S2.** Statistical Analysis. A. Friedman’s test was conducted to determine differences across AAV variants. The test was run separately for each cell type as well as total cells combined. Marmoset and cynomolgus macaque samples were both used in the analysis (n=8). Significant p-values < 0.05 are shown in red. S2.1 p-values resulting from a Friedman’s test using percent cells infected as the data points. Significant p-values < 0.05 are shown in red.

| Cell type | p-value |
| --- | --- |
| Rod | 2.22E-12 |
| Cone | 1.38E-08 |
| Horizontal Cell | 4.70E-12 |
| Off-Bipolar | 3.04E-12 |
| On-Bipolar | 1.26E-12 |
| Amacrine Cell | 3.81E-14 |
| Microglia | 1.14E-10 |
| Muller Glia | 5.73E-13 |
| Retinal Ganglion Cell | 4.50E-11 |
| Total Cells | 4.01E-15 |

**S2.2.** p-values resulting from a Friedman’s test using average transcripts per infected cell as the data points. Significant p-values < 0.05 are shown in red.

| Cell type | p-value |
| --- | --- |
| Rod | 5.45E-12 |
| Cone | 1.06E-08 |
| Horizontal Cell | 5.31E-12 |
| Off-Bipolar | 3.66E-12 |
| On-Bipolar | 1.74E-12 |
| Amacrine Cell | 1.89E-13 |
| Microglia | 1.36E-10 |
| Muller Glia | 3.15E-13 |
| Retinal Ganglion Cell | 1.91E-11 |

**Table S3.**
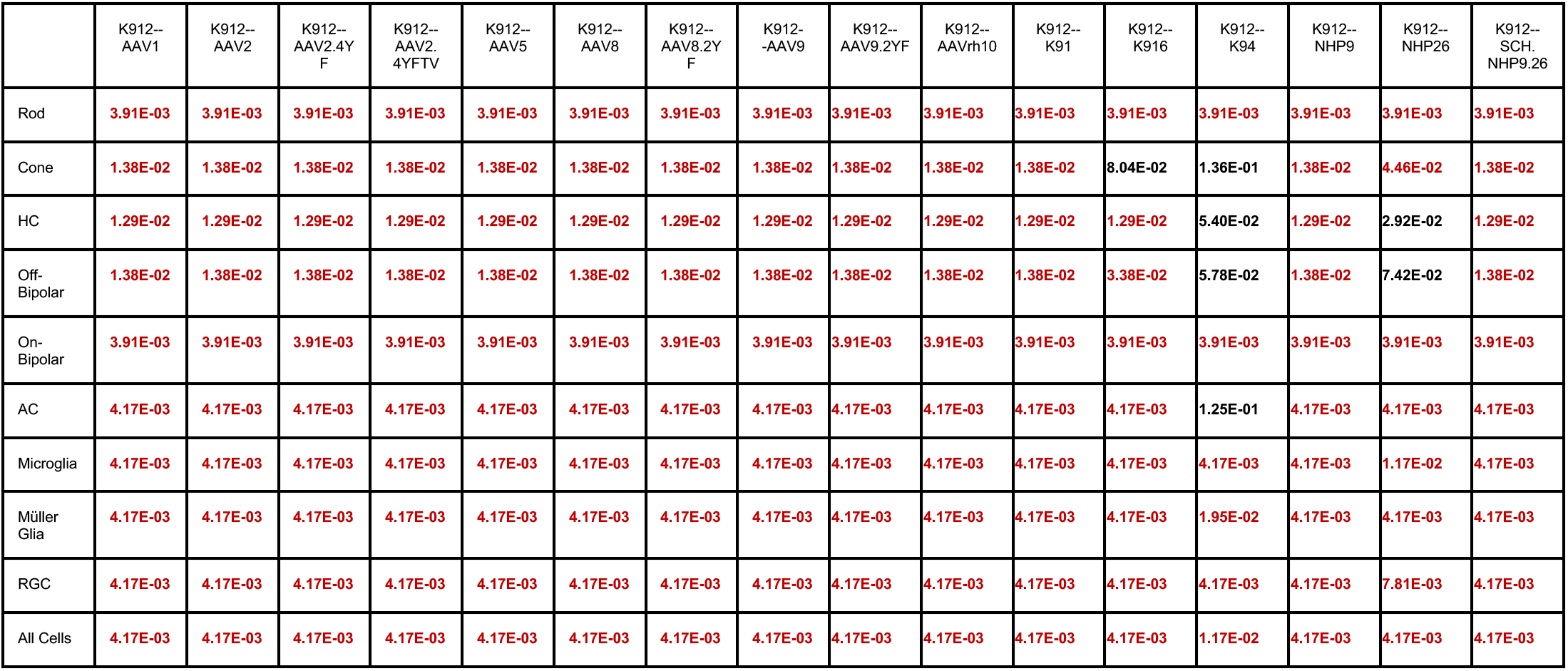
p-values from a Wilcoxon signed-rank test. A one-sided Wilcoxon signed-rank test was used for a pairwise comparison between K912 or NHP26 and the other variants. P-values were corrected using Benjamini-Hochberg correction method. Significant p-values < 0.05 are shown in red. S3.1 p-values resulting from a one-sided Wilcoxon signed-rank test using percent cells infected as the data points, comparing NHP12 and other variants.

**S3.2.** p-values resulting from a one-sided Wilcoxon signed-rank test using percent cells infected as the data points, comparing NHP26 and other variants.

|  | NHP26--<br>AAV1 | NHP26--<br>AAV2 | NHP26--<br>AAV2.4YF | NHP26--<br>AAV2.4YFTV | NHP26--<br>AAV5 | NHP26--<br>AAV8 | NHP26--<br>AAV8.2YF | NHP26--<br>AAV9 | NHP26--<br>AAV9.2YF | NHP26--<br>AAVrh10 | NHP26--<br>K91 | NHP26--<br>K912 | NHP26--<br>K16 | NHP26--<br>K94 | NHP26--<br>NHP9 | NHP26--<br>SCH.<br>NHP9.26 |
| --- | --- | --- | --- | --- | --- | --- | --- | --- | --- | --- | --- | --- | --- | --- | --- | --- |
| Rod | 4.81E-03 | 4.81E-03 | 4.81E-03 | 4.81E-03 | 4.81E-03 | 4.81E-03 | 4.81E-03 | 4.81E-03 | 4.81E-03 | 4.81E-03 | 4.81E-03 | 1 | 6.25E-02 | 6.71E-01 | 4.81E-03 | 4.81E-03 |
| Cone | 1.50E-02 | 4.64E-02 | 1.50E-02 | 1.50E-02 | 1.50E-02 | 1.50E-02 | 1.50E-02 | 1.50E-02 | 1.50E-02 | 1.50E-02 | 1.50E-02 | 9.73E-01 | 2.01E-01 | 6.05E-01 | 1.50E-02 | 1.50E-02 |
| HC | 4.81E-03 | 4.81E-03 | 4.81E-03 | 4.81E-03 | 4.81E-03 | 4.81E-03 | 4.81E-03 | 4.81E-03 | 4.81E-03 | 4.81E-03 | 4.81E-03 | 9.81E-01 | 8.30E-01 | 9.62E-01 | 4.81E-03 | 4.81E-03 |
| Off-Bipolar | 4.81E-03 | 4.81E-03 | 4.81E-03 | 4.81E-03 | 4.81E-03 | 4.81E-03 | 4.81E-03 | 4.81E-03 | 4.81E-03 | 4.81E-03 | 4.81E-03 | 9.45E-01 | 1.43E-01 | 5.62E-01 | 4.81E-03 | 4.81E-03 |
| On-Bipolar | 4.81E-03 | 4.81E-03 | 4.81E-03 | 4.81E-03 | 4.81E-03 | 4.81E-03 | 4.81E-03 | 4.81E-03 | 4.81E-03 | 4.81E-03 | 4.81E-03 | 1 | 1.79E-01 | 9.00E-01 | 4.81E-03 | 4.81E-03 |
| AC | 4.81E-03 | 4.81E-03 | 4.81E-03 | 4.81E-03 | 4.81E-03 | 4.81E-03 | 4.81E-03 | 4.81E-03 | 4.81E-03 | 4.81E-03 | 4.81E-03 | 1 | 3.12E-01 | 8.79E-01 | 4.81E-03 | 4.81E-03 |
| Microglia | 1.64E-02 | 1.64E-02 | 2.22E-02 | 2.22E-02 | 1.64E-02 | 1.64E-02 | 1.64E-02 | 1.64E-02 | 1.64E-02 | 1.64E-02 | 1.64E-02 | 9.92E-01 | 8.62E-02 | 8.93E-01 | 1.64E-02 | 1.64E-02 |
| Müller<br>Glia | 4.81E-03 | 4.81E-03 | 4.81E-03 | 4.81E-03 | 4.81E-03 | 4.81E-03 | 4.81E-03 | 4.81E-03 | 4.81E-03 | 4.81E-03 | 4.81E-03 | 1 | 3.12E-01 | 8.63E-01 | 4.81E-03 | 4.81E-03 |
| RGC | 4.81E-03 | 4.81E-03 | 4.81E-03 | 4.81E-03 | 4.81E-03 | 4.81E-03 | 4.81E-03 | 4.81E-03 | 4.81E-03 | 4.81E-03 | 4.81E-03 | 9.96E-01 | 1.79E-01 | 7.75E-01 | 4.81E-03 | 4.81E-03 |
| All Cells | 4.81E-03 | 4.81E-03 | 4.81E-03 | 4.81E-03 | 4.81E-03 | 4.81E-03 | 4.81E-03 | 4.81E-03 | 4.81E-03 | 4.81E-03 | 4.81E-03 | 1 | 2.19E-01 | 8.21E-01 | 4.81E-03 | 4.81E-03 |

**S3.3.** p-values resulting from a one-sided Wilcoxon signed-rank test using average transcripts per infected cell as the data points, comparing K912 and other variants.

|  | K912--<br>AAV1 | K912--<br>AAV2 | K912--<br>AAV2.4YF | K912--<br>AAV2.4YF<br>TV | K912--<br>AAV5 | K912--<br>AAV8 | K912--<br>AAV8.2YF | K912--<br>-AAV9 | K912--<br>AAV9.2YF | K912--<br>AAVrh10 | K912--<br>K91 | K912--<br>K916 | K912--<br>K94 | K912--<br>NHP9 | K912--<br>NHP26 |
| --- | --- | --- | --- | --- | --- | --- | --- | --- | --- | --- | --- | --- | --- | --- | --- |
| Rod | 4.17E-03 | 4.17E-03 | 4.17E-03 | 4.17E-03 | 4.17E-03 | 4.17E-03 | 4.17E-03 | 4.17E-03 | 4.17E-03 | 4.17E-03 | 4.17E-03 | 4.17E-03 | 1.17E-02 | 4.17E-03 | 4.17E-03 |
| Cone | 1.29E-02 | 1.29E-02 | 1.29E-02 | 1.29E-02 | 1.29E-02 | 1.29E-02 | 1.29E-02 | 1.29E-02 | 1.29E-02 | 1.29E-02 | 1.29E-02 | 1.29E-02 | 1.85E-02 | 1.29E-02 | 1.95E-02 |
| HC | 1.29E-02 | 1.29E-02 | 1.29E-02 | 1.29E-02 | 1.29E-02 | 1.29E-02 | 1.29E-02 | 1.29E-02 | 1.29E-02 | 1.29E-02 | 1.29E-02 | 1.45E-01 | 1.29E-02 | 1.29E-02 | 3.20E-01 |
| Off-Bipolar | 1.38E-02 | 1.38E-02 | 1.38E-02 | 1.38E-02 | 1.38E-02 | 1.38E-02 | 1.38E-02 | 1.38E-02 | 1.38E-02 | 1.38E-02 | 1.38E-02 | 1.85E-02 | 1.85E-02 | 1.38E-02 | 7.42E-02 |
| On-Bipolar | 4.46E-03 | 4.46E-03 | 4.46E-03 | 4.46E-03 | 4.46E-03 | 4.46E-03 | 4.46E-03 | 4.46E-03 | 4.46E-03 | 4.46E-03 | 4.46E-03 | 1.25E-02 | 1.91E-01 | 4.46E-03 | 4.46E-03 |
| AC | 4.81E-03 | 4.81E-03 | 4.81E-03 | 4.81E-03 | 4.81E-03 | 4.81E-03 | 4.81E-03 | 4.81E-03 | 4.81E-03 | 4.81E-03 | 4.81E-03 | 8.93E-03 | 7.42E-02 | 4.81E-03 | 4.17E-02 |
| Microglia | 4.17E-03 | 4.17E-03 | 4.17E-03 | 4.17E-03 | 4.17E-03 | 4.17E-03 | 4.17E-03 | 4.17E-03 | 4.17E-03 | 4.17E-03 | 4.17E-03 | 4.17E-03 | 4.17E-03 | 4.17E-03 | 2.73E-02 |
| Müller<br>Glia | 4.17E-03 | 4.17E-03 | 4.17E-03 | 4.17E-03 | 4.17E-03 | 4.17E-03 | 4.17E-03 | 4.17E-03 | 4.17E-03 | 4.17E-03 | 4.17E-03 | 4.17E-03 | 7.81E-03 | 4.17E-03 | 4.17E-03 |
| RGC | 3.91E-03 | 3.91E-03 | 3.91E-03 | 3.91E-03 | 3.91E-03 | 3.91E-03 | 3.91E-03 | 3.91E-03 | 3.91E-03 | 3.91E-03 | 3.91E-03 | 3.91E-03 | 3.91E-03 | 3.91E-03 | 3.91E-03 |

**S3.4.**
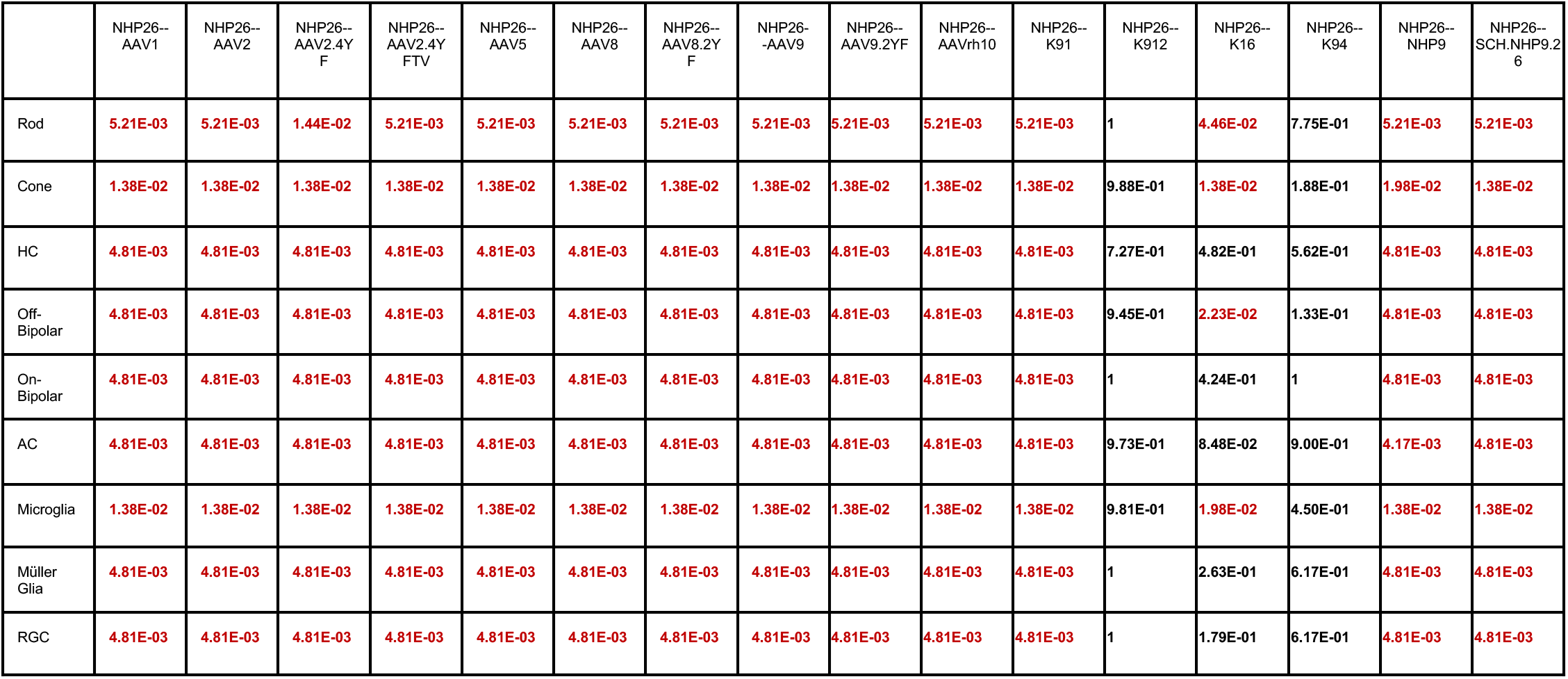
p-values resulting from a one-sided Wilcoxon signed-rank test using average transcripts per infected cell as the data points, comparing NHP26 and other variants.

**Table S4.**
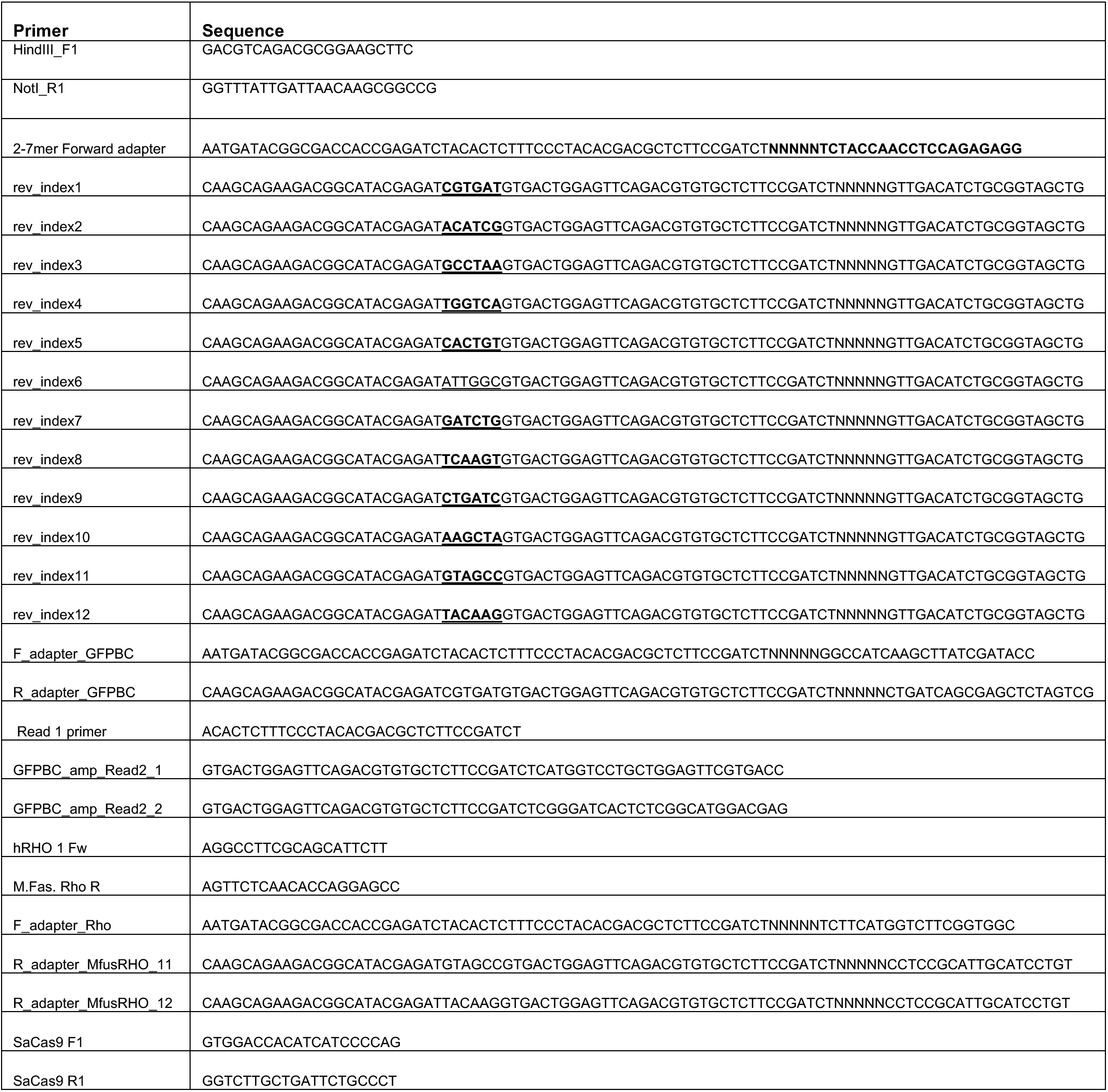
List of primers used in the study.

